# A convex optimization framework for gene-level tissue network estimation with missing data and its application in understanding disease architecture

**DOI:** 10.1101/2020.03.16.994020

**Authors:** Kushal K. Dey, Rahul Mazumder

## Abstract

Genes with correlated expression across individuals in multiple tissues are potentially informative for systemic genetic activity spanning these tissues. In this context, the tissue-level gene expression data across multiple subjects from the Genotype Tissue Expression (GTEx) Project is a valuable analytical resource. Unfortunately, the GTEx data is fraught with missing entries owing to subjects often contributing only a subset of tissues. In such a scenario, standard techniques of correlation matrix estimation with or without data imputation do not perform well. Here we propose `Robocov`, a novel convex optimization-based framework for robustly learning sparse covariance or inverse covariance matrices for missing data problems. `Robocov` produces more interpretable and less cluttered visual representation of correlation and causal structure in both simulation settings and GTEx data analysis. Simulation experiments also show that `Robocov` estimators have a lower false positive rate than competing approaches for missing data problems. Genes prioritized based on the average value of `Robocov` correlations or partial correlations across tissues are enriched for pathways related to systemic activities such as signaling pathways, heat stress factor, immune function and circadian clock. Furthermore, SNPs linked to these prioritized genes provide unique signal for blood-related traits; in comparison, no disease signal is observed for SNPs linked to genes prioritized by the standard correlation estimator. `Robocov` is an important stand-alone statistical tool for sparse correlation and causal network estimation for data with missing entries; and when applied to GTEx data, it provides insights into both genetic and autoimmune disease architectures.

## 1 Introduction

The gene expression data from nearly 50 tissues across more than 500 post-mortem donor individuals from Genotype Tissue Expression (GTEx) project has proved to be a valuable resource for understanding tissue-specific and tissue-shared genetic architecture^1,2,3,4,5,6^. Here we are interested in one specific aspect of tissue-shared gene regulation: the correlation and partial correlation in gene expression for different tissue pairs based on individual donor level data. A major challenge in this context is the extensive amount of missing entries in gene expression data—each donor contributes only a subset of tissues for sequencing. Common imputation based methods do not work well here as reported in ref.^7^, owing to stringent assumptions about missing entries being close to some central tendency (median) or adhering to some low-dimensional representation of the observed entries^8,9,10^. Popular shrinkage and/or sparse correlation or partial correlation estimators such as *corpcor*^11,12,13^, GLASSO^14^ or CLIME^15^ are not designed for data with missing values.

A recently proposed approach, `CorShrink`^7^, co-authored by one of the authors (Dey), does account for this missing information through adaptive shrinkage^16^ of correlations. `CorShrink` does not guarantee a positive semidefinite (PSD) matrix as part of its EM-based framework, and necessitates a post-hoc modification to ensure a PSD correlation matrix. Also, `CorShrink` does not extend to conditional graph or partial correlation estimation. Here, we propose a new approach based on convex optimization, called `Robocov`—this applies for both covariance and inverse covariance matrix estimation in the presence of missing data under the following regularization principles:

- the covariance matrix is sparse (i.e., has a few nonzero entries)
- inverse covariance matrix is sparse.

`Robocov` does not *impute* missing values per-se^1^-it directly estimates the covariance or inverse covariance matrices in the presence of missing values. To handle missing values, we consider a loss function that depends upon the pairwise covariance terms (computed based on the observed samples) but incorporates an adjustment to guard against our lack of knowledge regarding the missing observations. For inverse covariance estimation, `Robocov` uses a robust optimization based approach^18,19^ that accounts for the uncertainty in estimating the pairwise sample covariance terms (due to the presence of missing values). Interestingly, both lead to convex optimization formulations that are amenable to modern optimization techniques^20^—they are scalable to moderate-large scale instances; and unlike conventional EM methods (that lead to highly nonconvex optimization tasks), our estimators are guaranteed to reach the optimal solution of the optimization formulations that define the `Robocov` estimators.

Our simulation experiments suggest that `Robocov` estimators for correlation and partial correlation matrices have a lower false positive rate compared to competing approaches when data has missing entries. When applied to the GTEx gene expression data comprising of ~ 70% missing data, `Robocov` produces less cluttered and highly interpretable visualization of correlation and conditional graph architecture, compared to standard approaches.

From a biological perspective, a gene with high correlation in expression across many tissue pairs is potentially reflective of systemic biological processes affecting many tissues and organs. To this end, we prioritize genes based on the average `Robocov` estimated correlation (partial correlation) across all tissue-pairs; we call them `Robospan` (`pRobospan`) genes. A pathway enrichment analysis of `Robospan` (`pRobospan`) genes showed enrichment in systemic functional pathways such as interferon signaling, heat stress factors, circadian clock and more importantly, the immune system. Subsequently we generated SNP level annotations for SNPs linked to `Robospan` (`pRobospan`) genes and tested for autoimmune disease informativeness by applying Stratified LD-score regression (S-LDSC) to 11 common blood-related traits (5 autoimmune diseases and 6 blood cell traits; average *N*=306K), conditional on a broad set of coding, conserved, regulatory and LD-related annotations. `Robospan` and `pRobospan` genes showed high disease informativeness for blood and autoimmune diseases and traits; in comparison; the analogously defined `Corspan` genes defined using the standard correlation estimator was non-informative. This highlights the biological and disease-level significance of our work.

In Section 2, we present an overview of methods and the underlying optimization framework for `Robocov`. Section 3 presents the simulation results and the application of `Robocov` to GTEx gene expression data and the downstream application of `Robocov` in understanding the autoimmune disease architecture. Finally, Section 4 presents an overall summary and future directions.

## 2 Methods and Materials

Let *X_N×P_* be a data matrix with *N* samples and *P* features, where some of the entries *X_np_* may be missing, denoted here by NA. We let *X^f^* denote the fully-observed version of the partially-observed data matrix^2^ *X*. We further assume that the fully observed data vectors 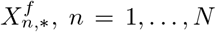 are independent and follow a Multivariate Normal distribution:

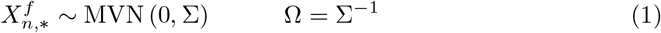

where Σ*_P×P_* and Ω*_P×P_* denote the model covariance and the inverse covariance matrices respectively. Based on the observed entries, we obtain a matrix 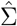 of pairwise covariances such that for all *i*, *j* ∈ {1,…, *P*}:

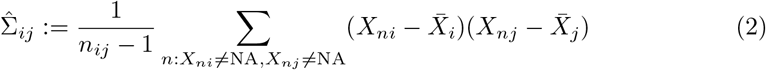

where, 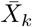 denotes the sample mean of feature *k* based on the observed entries; and *n_ij_* is the number of samples *n* with non-missing entries in both features *i* and *j*:

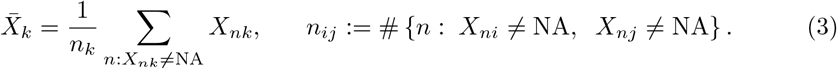

Here *n_k_* denotes the number of observed samples (i.e., not missing) for feature *k*. For our analysis, we will assume^3^ that *n_ij_* > 2 for all *i, j*. We note that the matrix of all pairwise covariance terms: 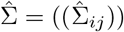, as defined in (2), need not be positive semidefinite due to the presence of missing values in the data matrix.

### 2.1 Robocov covariance estimator

We first present the `Robocov` covariance matrix estimator—this leads to an estimate of Σ via the following regularized criterion:

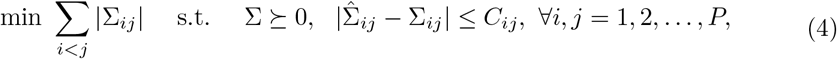

where Σ is the optimization variable and 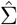 and *C_ij_* s are problem data. Note that Problem (4) minimizes a convex penalty function subject to convex constraints — the optimization variable Σ is positive-semidefinite (denoted as Σ ≽ 0). Hence (4) is a convex semidefinite optimization problem^20^; and can be solved efficiently by modern semidefinite optimization algorithms for moderately large instances (e.g, *P* ~ 1000) using (for example) the SCS solver in CVX software^20,21,22,23^. The objective function in (4) minimizes the *ℓ*_1_-norm on the entries of Σ; and induces sparsity in the solution^24^. The constraint 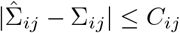 for all *i,j* is the data-fidelity term — it constrains the entries of the estimated covariance matrix (i.e., Σ*_ij_*) to be close to the sample covariance 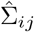—that is, 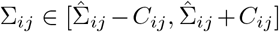. Here, *C_ij_* controls the amount by which Σ*_ij_* can differ from the sample version 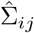. We compute *C_ij_* based on the Fisher’s Z-Score^25,26^(for a complete derivation see Supplementary Note):

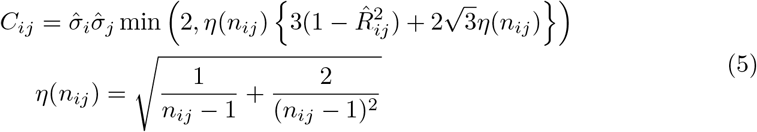

where 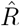 is the pairwise sample correlation matrix derived from 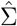 i.e.,

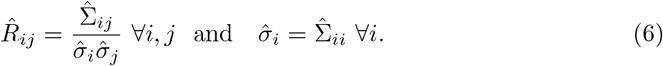

Note that criterion (4) can be perceived as a special case of a more general regularized optimization problem

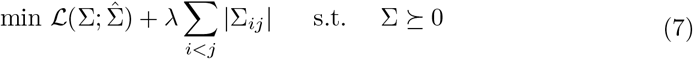

where, 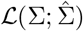 is the data-fidelity term or loss function measuring the proximity of Σ to 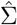; the second term represents the regularization on Σ; and λ is a tuning parameter that controls the trade-off between data-fidelity and regularization. We can choose 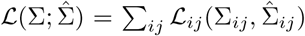 with 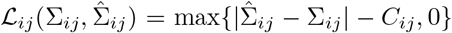 for all *i,j*. This leads to a regularized convex optimization problem of the form:

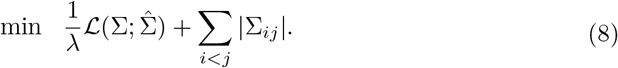

In the limiting case, λ → 0+ i.e., 1/λ → ∞, estimator obtained from Problem (8) will reduce to the estimator available from (4). This is because, for sufficiently large values of 1/λ, an optimal solution to (8) will lead to a zero loss—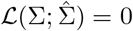 which implies that 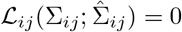 for all *i, j* — these are the data-fidelity constraints in (4).

In summary, we note that our proposed `Robocov` estimator does not impute missing values per-se — it directly leads to an estimate for the covariance matrix Σ while taking into account the presence of missing-values in the data matrix.

Other choices of the loss function 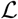 are also possible — we present one such candidate in the Supplementary Note that is also derived from the Fisher’s Z-Score setup (which was key to deriving (4)). In practice, we found that many of these estimators lead to similar results on real datasets—therefore, in this paper, we focus our attention on the basic estimator (4).

#### From covariance to correlation estimates

Formulation (4) delivers an estimate of the covariance matrix. To obtain a correlation matrix estimate, one can modify (4) to deliver a correlation matrix instead of a covariance matrix:

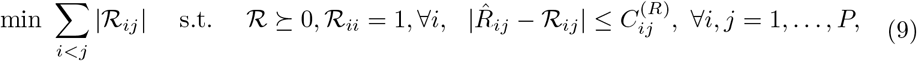

where 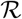 is the optimization variable, 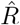 is defined in (6) and 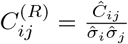. The derivation of *C*^(*R*)^ follows from the derivation of *C* and appears in the Supplementary Note.

A simple alternative approach to obtain the `Robocov` correlation matrix estimate, is to re-scale the `Robocov` covariance estimator (obtained from Problem (4)) to a correlation matrix.

### 2.2 Robocov inverse covariance estimator

Section 2.1 discusses a convex optimization-based estimator for the covariance matrix (Σ), here we present a method to estimate the inverse covariance matrix (Ω) and subsequently the partial correlation matrix. We present a regularized likelihood framework to estimate Ω under a sparsity constraint. An appealing aspect of our estimator is that our optimization criterion is convex in Ω (and not Σ which was the case in Section 2.1).

Our estimator builds upon the popular *ℓ*_1_-regularized Gaussian likelihood framework (aka graphical lasso or GLASSO^14,27,28^) for the fully observed case, and adapts it to address missing values. We recall the GLASSO procedure which minimizes an *ℓ*_1_-norm regularized negative log-likelihood criterion (fully observed case) given by:

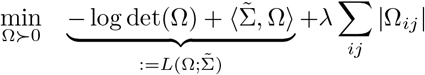

where, 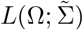 is the negative log-likelihood (ignoring irrelevant constants) for the model (1), 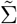 is the fully observed sample covariance matrix and λ ≥ 0 is the regularization parameter.

Replacing 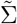 by the observed matrix 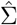 in 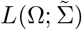 is problematic due to the error in estimating the pairwise covariances arising from the missing values (different cell entries of the sample covariance matrix involve different effective sample sizes *n_ij_*s leading to varying accuracies in estimating 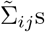). To account for this uncertainty, we use ideas from robust optimization^18,19^—to the best of our knowledge, this approach has not been used earlier in the context of sparse inverse covariance estimation (in the presence of missing values). Our robust optimization approach minimizes the worst-case loss arising from the errors in estimating the cell entries 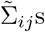. This leads to a min-max optimization problem of the form:

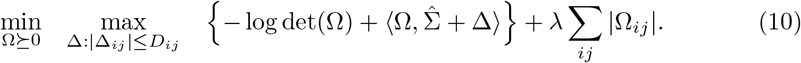

As (10) involves minimization of a pointwise maximum (over Δ) of convex functions 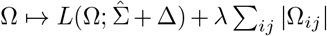, Problem (10) is convex^20^ in Ω. Convexity ensures that a global minimum to the problem can be obtained reliably—making our approach different from traditional missing data techniques based on the EM algorithm^17^ that often lead to complex nonconvex optimization tasks with multiple local solutions.

In words, the inner maximization over Δ in Problem (10) gives the largest (or worst-case) value of the negative log-likelihood—max_Δ_ 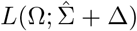 where, Δ captures the uncertainty involved in estimating the entries of the sample covariance matrix 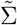 due to the presence of missing values. The outer minimization problem (wrt Ω) considers the minimum of the *adjusted* worst-case loss function 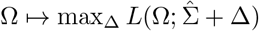, in addition to an *ℓ*_1_-penalization on Ω that encourages a sparse estimate of Ω.

The so-called uncertainty set^19^ in Δ is given by: |Δ*_j_*| ≤ *D_ij_* (for all *i, j*) where, the upper bound *D_ij_* arises from a probability computation using the Fisher’s Z-score criterion (see Supplementary Note for additional details):

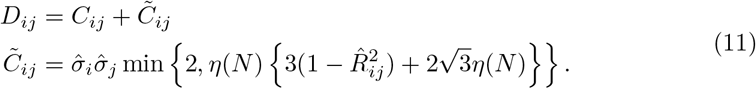

To provide some intuition about (11), the value of the error *D_ij_* will be large if *n_ij_* is small, and will be equal to zero when *n_ij_* = *n* (with no missing entries).

The seemingly complicated min-max optimization problem in (10) reduces to a cousin of the GLASSO criterion (See Supplementary Note for details) — we use a weighted version of the *ℓ*_1_-norm penalty on Ω:

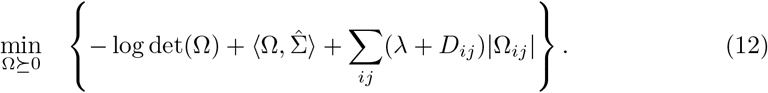

Problem (12) is a nonlinear semidefinite optimization problem in Ω—and the constraint Ω ≽ 0 leads to a positive semidefinite inverse covariance matri^4^. Problem (12) uses a weighted *ℓ*_1_-norm on Ω where the penalty weights are adjusted to account for the uncertainty due to the presence of missing values. Note that the penalty parameter λ accounts for the sparsity in Ω arising from our prior sparsity assumption on Ω—the overall penalty weight for the (*i,j*)-th entry, (λ + *D_ij_*) adds further regularization due to the presence of missing values. In particular, if there is no missing value, then *D_ij_* = 0 and (12) will reduce to the GLASSO criterion. If *n_ij_* is small, then the value of *D_ij_* will be large — therefore, we will place a higher weight on the term |Ω*_ij_*| to shrink it towards zero.

Note that, as in Section 2.1, the `Robocov` inverse covariance estimator, bypasses the task of imputing the missing values. Our main goal is to directly estimate Ω from a partially observed data-matrix *X*. In this way, we can potentially mitigate the limitations of a sub-optimal imputation procedure. See Section 3 for an empirical validation.

Criterion (12) leads to an inverse covariance estimator — we use the solution Ω from Problem (12) to define a partial correlation estimator 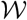 as follows

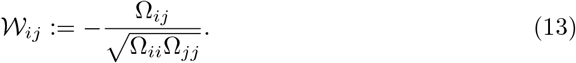

Both the optimization problems (4) and (12) were solved using R implementation of the CVX software^22,23^. This was sufficient for the problem-scales we are dealing with —for larger instances, specialized algorithms (e.g., based on first order methods)^24,29,30^ may be necessary.

In all our subsequent analysis and numerical results, we use the `Robocov` correlation estimator 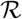 (see Problem (9)) and partial correlation estimator 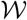 (13).

## 3 Results

### Simulation Experiments: Synthetic and Real Data

We applied `Robocov` on simulated multivariate normal data from three population correlation structure models (hub, Toeplitz and 1-band precision matrix) with *N* samples, *P* features and *π* proportion of missing entries randomly distributed throughout the data matrix (see Supplementary Note for details). For ease of interpretation, the features have unit variance under all three models, implying that the population covariance matrix is the same as the correlation matrix.

Figure 1 shows results for all three model-settings with *N* = 500, *P* = 50, *π* = 0.5. For every setting, `Robocov` generated a sparse estimate of the population correlation 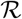 (Section 2.1) or population partial correlation 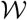 (Section 2.2). The `Robocov` correlation estimator captured the population structure effectively for all three models, while the standard pairwise sample correlation estimator (based on the observed entries) showed comparatively poor performance (Figure S1). The `Robocov` partial correlation estimator accurately captured the causal structure in the hub and 1-band precision matrix models; for the Toeplitz matrix, it captured the high partial correlation band immediately flanking the diagonal but failed to capture the other alternating positive and negative low correlation bands (Figure 1).

**Figure 1.**
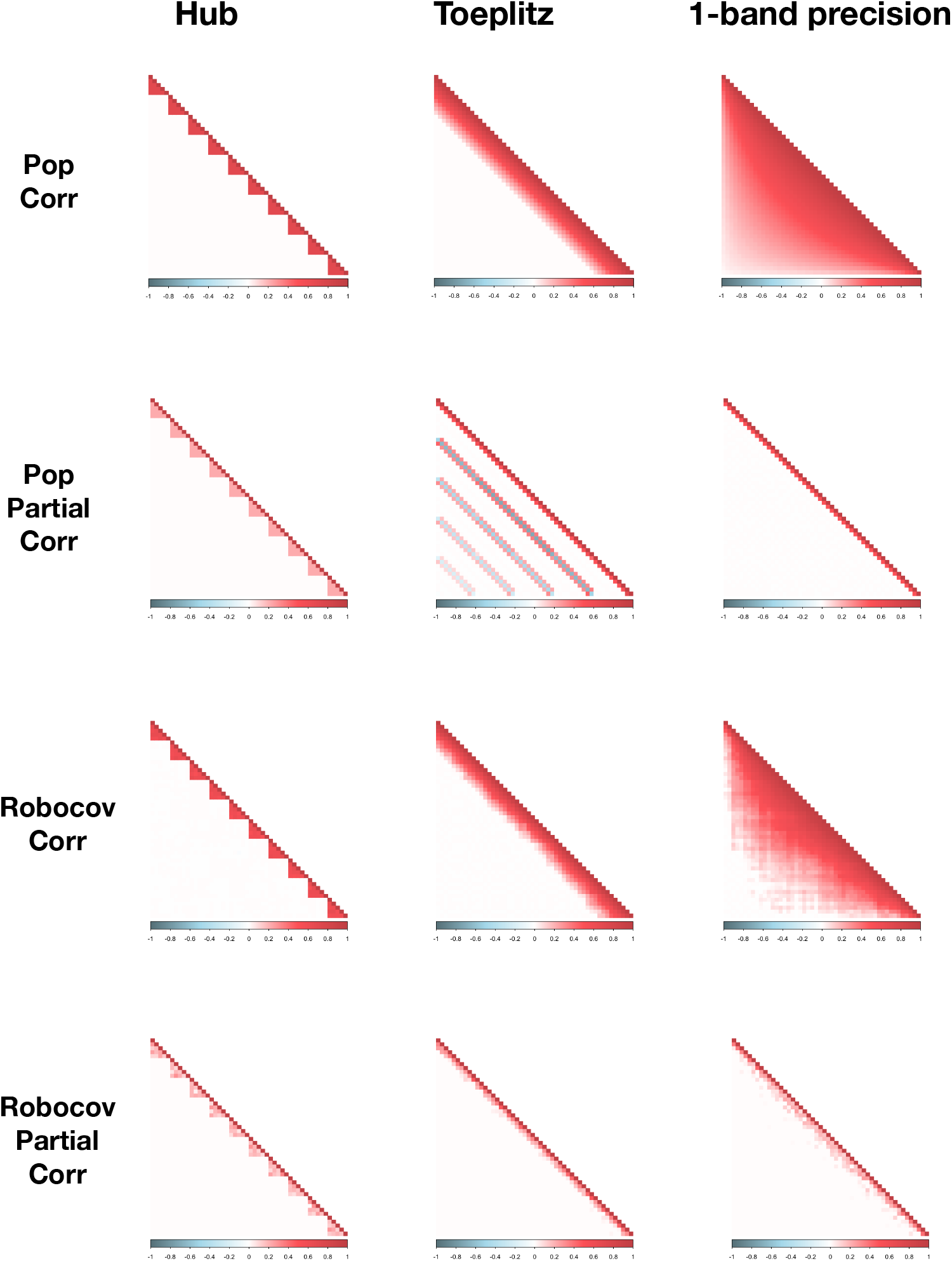
Simulation results of applying Robocov correlation and partial correlation estimators on Hub, Toeplitz and 1-band precision correlation structures: We applied `Robocov` correlation and partial correlation estimators on data generated from Hub, Toeplitz or 1-band precision matrix based population models (see Simulation settings in Supplementary Note) with *N* = 500 samples, *P* = 50 features and *π* = 0.5 proportion of missing data. We present the population correlation matrix in the first row, population partial correlation matrix in second row, `Robocov` correlation matrix in third row and `Robocov` partial correlation matrix in last row. The tuning parameter for `Robocov` correlation matrix (and partial correlation matrix) estimation was determined by cross-validation.

In view of the biological problem of interest, the hub structured population correlation model is perhaps the most interesting—recent work^7^ has shown hub-like patterns in expression correlation across tissue pairs for most genes. To this end, we applied `Robocov` on simulated data for hub population correlation matrix structure for different settings of *N, P* and *π* (see Supplementary Note for details). Two metrics of particular interest were the false positive rate (FPR) and the false negative rate (FNR) as defined in Supplementary Note. Using these metrics, we compared the `Robocov` correlation estimator with both the pairwise sample correlation estimator and the recently proposed adaptive shrinkage based approach, `CorShrink`. Across different (*N, P*, *π*)-settings, the `Robocov` correlation estimator had lower FPR than `CorShrink`. In comparison, for data with a large number of missing entries (i.e., high *π*), FNR for `Robocov` was worse compared to `CorShrink` (Table 1). We did not compare against other shrinkage-based correlation estimators such as *PDSCE*^31^ and *corpcor*^32,13^ as (i) they do not account for missing entries in the data and (ii) even in the fully observed case (i.e., no missing values) earlier work^7^ has shown that these methods are outperformed by `CorShrink` (see Figure 4 from ref.^7^).

**Table 1.** Simulation results of correlation and partial correlation estimators: Hub population structure model. We report metrics to compare (i) the `Robocov` correlation estimator (*Cor*) against `CorShrink` and the standard pairwise sample correlation estimator; and (ii) the `Robocov` partial correlation estimator (*P.Cor*) against estimators available from GLASSO and CLIME. Here, data is generated from a hub-structured population covariance matrix with different choices of *N* (number of samples), *P* (number of features) and different degrees of missing entries in the data (the fraction *π* of missing data varies from 0% to 50%). The three metrics are FP2 (False Positive 2-norm), FPR (False Positive Rate) and FNR (False Negative Rate). See Supplementary Note for the details of the metrics. Results are averaged over 50 replications from the same model. For all three partial correlation estimators: `Robocov` partial correlation, GLASSO and CLIME; the optimal sparsity inducing parameter λ was chosen by cross-validation. See Simulation settings under Supplementary Note for further details on the simulation model.

| Hub: N = 50, P=50 |  |  |  |  |  |  |  |  |  |  |
| --- | --- | --- | --- | --- | --- | --- | --- | --- | --- | --- |
| Type | Method | $\pi=0$ | | | $\pi=0.25$ | | | $\pi=0.5$ | | |
|  |  | FP2 | FPR | FNR | FP2 | FPR | FNR | FP2 | FPR | FNR |
| Cor | Robocov | 0.047 | 0 | 0 | 0.14 | 0 | 0.14 | 0.26 | 0 | 0.19 |
|  | CorShrink | 1.4 | 0.01 | 0 | 2.2 | 0.04 | 0.03 | 4 | 0.07 | 0.09 |
|  | Standard | 6.7 | 0.24 | 0 | 8.8 | 0.30 | 0 | 15 | 0.28 | 0 |
| P.Cor | Robocov | 0.08 | 0 | 0.07 | 0.27 | 0.01 | 0.13 | 0.47 | 0 | 0.09 |
|  | GLASSO | 0.12 | 0 | 0.15 | 0.29 | 0.01 | 0.15 | 0.59 | 0.02 | 0.12 |
|  | CLIME | 1.47 | 0.09 | 0.07 | 1.37 | 0.07 | 0.08 | 1.29 | 0.08 | 0.07 |
| Hub: N = 100, P=50 |  |  |  |  |  |  |  |  |  |  |
| Type | Method | $\pi=0$ | | | $\pi=0.25$ | | | $\pi=0.5$ | | |
|  |  | FP2 | FPR | FNR | FP2 | FPR | FNR | FP2 | FPR | FNR |
| Cor | Robocov | 0.046 | 0 | 0 | 0.062 | 0 | 0 | 0.18 | 0 | 0.15 |
|  | CorShrink | 0.9 | 0 | 0 | 1.3 | 0.02 | 0 | 2.9 | 0.03 | 0.01 |
|  | Standard | 4.8 | 0.17 | 0 | 6.2 | 0.20 | 0 | 10 | 0.31 | 0 |
| P.Cor | Robocov | 0.23 | 0 | 0.06 | 0.21 | 0 | 0.09 | 0.18 | 0.03 | 0.11 |
|  | GLASSO | 0.11 | 0 | 0.16 | 0.23 | 0 | 0.22 | 0.29 | 0.01 | 0.24 |
|  | CLIME | 1.83 | 0.12 | 0.08 | 1.77 | 0.14 | 0.09 | 1.78 | 0.16 | 0.11 |
| Hub: N = 500, P=50 |  |  |  |  |  |  |  |  |  |  |
| Type | Method | $\pi=0$ | | | $\pi=0.25$ | | | $\pi=0.5$ | | |
|  |  | FP2 | FPR | FNR | FP2 | FPR | FNR | FP2 | FPR | FNR |
| Cor | Robocov | 0.028 | 0 | 0 | 0.012 | 0 | 0 | 0.077 | 0 | 0 |
|  | CorShrink | 0.21 | 0 | 0 | 0.32 | 0 | 0 | 0.83 | 0 | 0 |
|  | Standard | 2.1 | 0.01 | 0 | 2.8 | 0.05 | 0 | 4.4 | 0.14 | 0 |
| P.Cor | Robocov | 0.12 | 0 | 0.11 | 0.16 | 0 | 0.12 | 0.11 | 0 | 0.14 |
|  | GLASSO | 0.16 | 0 | 0.19 | 0.29 | 0 | 0.20 | 0.19 | 0.02 | 0.20 |
|  | CLIME | 2.11 | 0.11 | 0.16 | 1.99 | 0.14 | 0.18 | 2.04 | 0.15 | 0.17 |

Next, we assess the performance of the `Robocov` partial correlation estimator for the same simulation settings (Table 1). We are not aware of a sparse conditional graph or partial correlation estimation method that directly takes into account missing entries. Nevertheless, we compare the `Robocov` partial correlation estimator with (i) GLASSO on the pairwise sample correlation estimator 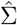 and (ii) CLIME on an imputed data matrix where, the imputation is performed using SoftImpute^9^. Comparisons are made in terms of FPR and FNR. In the presence of missing data, `Robocov` partial correlation estimator showed better FPR and FNR compared to both GLASSO and CLIME-based estimators (Table 1). CLIME is found to under-perform in our experiments—this may largely be due to the error arising from the imputation step (Table 1).

Next, we evaluated the predictive performance of `Robocov` with pairwise sample correlation estimator and `CorShrink`. We considered the GTEx gene expression data for a particular gene (ARHGAP30) across 544 donors and 53 tissues with close to 70% missing data owing to subjects contributing only a small fraction of tissues. We split the individual by tissue data for the gene into two equal groups and compared the estimated correlation matrix (we used different estimators: `Robocov`, `CorShrink` and pairwise sample correlation matrix) computed on one half of the individuals with the pairwise sample correlation matrix computed from the other half. Both `Robocov` and `CorShrink` estimators considerably outperformed the pairwise sample correlation estimator, with `CorShrink` having slightly better predictive accuracy (Figure S2 and Table S1). Due to the similar predictive performances of `Robocov` and `CorShrink`, the former may be preferable as it results in sparse estimates, leading to better interpretability.

An an alternative to `Robocov`, we may consider an estimator obtained by first imputing the missing entries in the data matrix and then estimating the correlation or partial correlation matrix for the complete data. For the same ARHGAP30 gene, we performed imputation by either a low rank factorization (SoftImpute^9^, with or without scaling) or a median based approach (replacing the missing entries of a feature by the median value of the observed entries). The correlation matrix obtained by SoftImpute (both with and without scaling) showed artificial high negative and positive correlation sweeps between brain and non-brain tissues that were not observed in the pairwise correlation matrix (Figure S3). One possible explanation of this is that the data matrices in our case do not seem to have a low rank representation based on eigenvalue analysis (Figure S4). The median based imputation method on the other hand, is prone to showing false positives—for example, we see a high correlation between Fallopian tube and Cervix-Ectocervix, which is a consequence of only 3 individuals contributing both the tissues (Figure S3). `Robocov` can effectively get rid of these edge cases and generate sparser and more robust results compared to these imputation based approaches.

Based on our simulation studies, we conclude that the `Robocov` correlation estimator has a lower FPR than both the standard pairwise sample correlation estimator and `CorShrink`. In terms of predictive performance, `Robocov` does better than the standard estimator and is comparable to `CorShrink`. We also observe that for data with a large number of missing entries and no obvious low rank representation as in case of the GTEx gene expression data, imputation based approaches are sub-optimal and `Robocov` would be the preferred option in such a scenario. The `Robocov` partial correlation estimator, on the other hand, showed better performance both in terms of FPR and FNR compared to other competing methods such as GLASSO and CLIME, especially when the proportion of missing entries in the data matrix is high.

### Gene Expression correlation analysis across tissue pairs

We applied `Robocov` to each of 16,069 cis-genes (genes with at least one significant cis-eQTL) from the GTEx v6 project^5,4^ (see URLs for gene list). For each gene, the data matrix had 544 rows (post-mortem donors), 53 columns (tissues) and comprised of ~ 70% missing entries. The median `Robocov` correlation estimator showed weak hub-like association across 13 Brain tissues, 3 Artery tissues, 3 Esophagus tissues, 2 Heart tissues and 2 Skin tissues (Figure S5). Figure 2 presents a visual comparison of `Robocov` correlation and partial correlation estimators with standard pairwise sample correlation matrix for two example genes (ARHGAP30 and GSTM1)—the `Robocov` correlation estimator is sparse and visually less cluttered than the standard approach. The `Robocov` correlation structure across tissue pairs varied from one gene to another: some genes showed high correlation across all tissues (e.g. HBB, RPL9), some showed little correlation across tissues (e.g. NCCRP1), some showed high intra-Brain correlation but relatively low inter-Brain correlation (e.g. ARHGAP30) (Figure 3, Figure S6 and Figure S2). Furthermore, two genes with similar correlation profiles may have very distinct expression profiles. For example, HBB and RPL9 showed high correlation across all tissue pairs, but they were distinct in their tissue-specific expression profiles. HBB showed high expression in Whole Blood relative to other tissues, while RPL9 had a more uniform expression profile across tissues (Figure 3). A similar pattern was observed also for two genes with negligible correlation across tissues, NCCRP1 and RPL21P11 (Figure S6).

**Figure 2.**
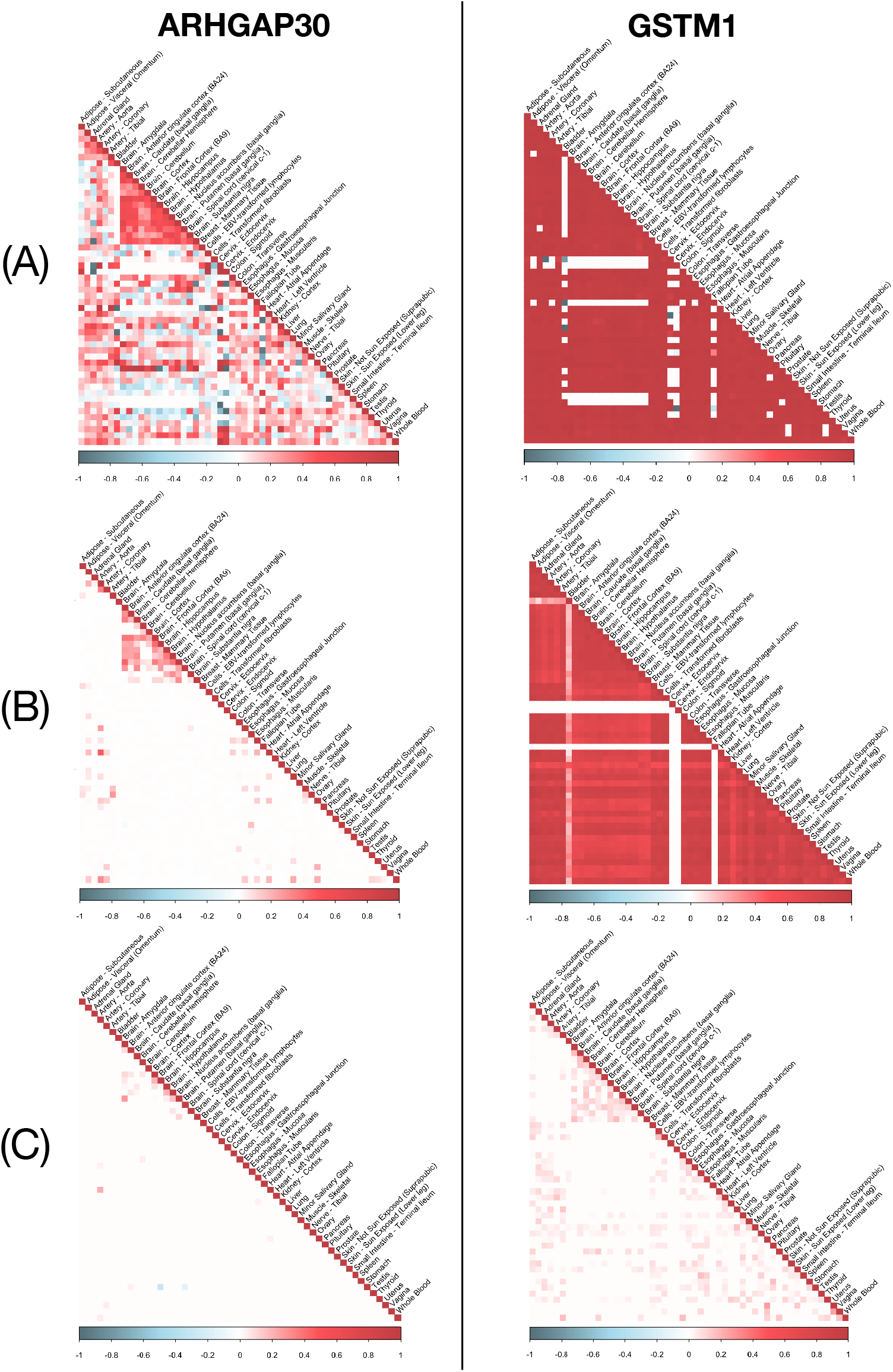
Illustrative examples of pairwise sample correlation estimator, Robocov correlation and partial correlation estimators for 2 genes: (Left column) **ARHGAP30** gene and (Right column) **GSTM1** gene. Each column shows the (A) pairwise sample correlation estimator, (B) `Robocov` correlation estimator and (C) partial correlation estimator stacked from top to bottom.

**Figure 3.**
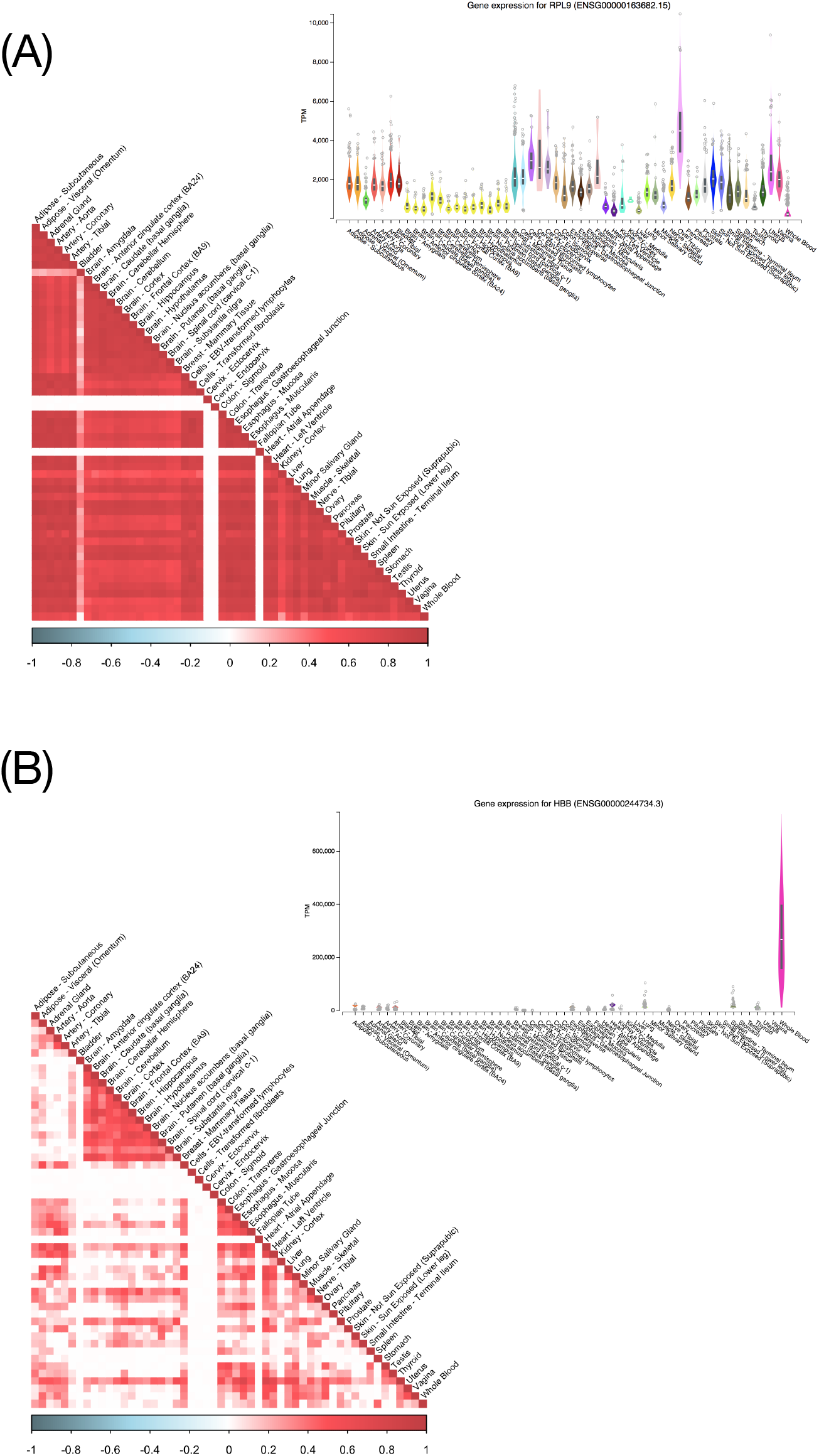
Examples of genes with high average Robocov correlation across all tissue pairs but with distinct expression profiles: (A) **RPL9** gene has uniformly high TPM (transcripts per million) values across most tissues (inset picture). (B) **HBB** shows high expression specifically in Whole Blood (inset picture). The expression profile plots for the genes have been fetched from the GTEx Portal (https://gtexportal.org/home/).

Next, we assign to each gene, a prioritizing score defined by the average value of `Robocov` correlation (*Robospan-score*) or partial correlation (*pRobospan-score*) across all tissue pairs. Similarly, we also computed the average value of the pairwise sample correlation (*Corspan-score*) across tissues. Then we tested and compared these three gene prioritizations based on how well they capture functional and disease relevant architecture. None of the three scores showed significant enrichment in 3,804 housekeeping genes^33^ (0.84x, 0.72x and 0.4x for `Robospan`-score, Corspan-score and `pRobospan`-score respectively). We compared these 3 gene scores with constraint-based metric of gene essentiality such as the absence of loss-of-function(LoF) variants (pLI^34^ and s_het^35^). For each of the 50 quantile bins of pLI and s_het, we computed the median of each of these scores; and compared with the mid-value of the quantile bin. We observed a slight negative trend in all 3 scores with increasing quantile bins of both pLI (*r* = −0.07 for Robospan-score, −0.13 for pRobospan-score and −0.13 for Corspan-score) and s_het (*r* = −0.05 for Robospan-score, −0.10 for pRobospan-score and −0.08 for Corspan-score) (Figure S7). One possible explanation may be that genes with highly correlated expression across all tissues may be driven by tissue-shared regulation machinery which imposes lower selective constraints on these genes.

The top 10% genes from each of the three gene prioritizing scores were used to define gene sets; we call them `Robospan`, `pRobospan` and `Corspan` genes. We performed a pathway enrichment analysis^36,37^ of these gene sets; the top 5 enriched pathways included immune system, interferon signaling, heat stress factor (Table S2). The magnitude of enrichment was stronger for `Robospan` and `pRobospan` genes compared to Corspan genes. Though not among the top 5 pathways, other interesting significant pathways included different signaling pathways (interleukin mediated signaling, VEGFA-VEGFR2, NFkB signaling) and Circadian clock related pathways (see URLs). The enrichment of immune related pathways was further backed by high enrichment of these genes in top 10% specifically expressed genes in Whole Blood (SEG-Blood^38^) (`Robospan`: 1.48x, `pRobospan`: 2.50x, `Corspan`: 1.45x). One possible conjecture may be that this enrichment is an artifact caused by contamination of blood with GTEx tissue samples. This, however, is countered by examples of genes that have high correlation across all tissues but expression-wise, are specific to tissues that are not Whole Blood (Figure S8). We also see examples of specifically expressed genes in Whole Blood that have low Robospan-score (Figure S9). The other possible reason may be biological; some highly expressed genes in blood may carry out important systemic functional activity across different tissues (cell-cell signaling, transport of substances, immune response) and therefore show high correlation across tissues.

We conclude that `Robocov` produces less visually cluttered representation of correlation and partial correlation structure of gene expression across tissue pairs for individual genes. We also show that genes with high average `Robocov` correlation or partial correlation across tissue pairs tend to have lower selection constraint and are not enriched for housekeeping genes. The top genes with highest average `Robocov` correlation or partial correlation across tissues are enriched for immune related functionality among other systemic pathways such as heat stress factors, circadian clock etc. This is further backed by enrichment of `Robospan` and `pRobospan` genes with specifically expressed genes in Blood.

### Heritability analysis of blood-related traits

The strong connection of `Robospan`, `pRobospan` and `Corspan` genes with blood-related genes and immune related pathways, as reported in the previous subsection, prompted us to test whether these genes are uniquely informative for blood-related complex diseases and traits.

For each gene set, we define SNP-level annotations to test for disease heritability. We define an *annotation* as an assignment of a numeric value to each SNP with minor allele count ≥5 in a 1000 Genomes Project European reference panel^39,40^. For each gene set X, we generate two binary SNP-level annotations – we assign a value of 1 to a SNP if it lies within 5kb or 100kb window upstream and downstream of a gene in the gene set and 0 otherwise; this strategy has been used in several previous works^38,41,42^.

We assessed the informativeness of the 6 SNP annotations (2 SNP annotations per gene set) for disease heritability by applying stratified LD score regression (S-LDSC)^40^ conditional on 86 baseline annotations comprising of coding, conserved, epigenomic and LD related annotations (this is called the baseline-LD model; here we use version 2.1^43^). S-LDSC results were meta-analyzed across 11 relatively independent blood-related traits (5 autoimmune diseases and 6 blood traits (Table S3). We considered two S-LDSC metrics for comparison: enrichment and standardized effect size (*τ*^⋆^) (see Supplementary Note for details). Enrichment is defined as the proportion of heritability explained by SNPs in an annotation divided by the proportion of SNPs in the annotation^40^. Standardized effect size (*τ*^⋆^) is defined as the proportionate change in per-SNP heritability associated with a 1 standard deviation increase in the value of the annotation, conditional on other annotations included in the model^43,44^; unlike enrichment, *τ*^⋆^ quantifies effects that are unique to the focal annotation. Here we primarily use *τ*^⋆^ as a metric for disease informativeness like in several previous works^7,38,41,44,45^.

All 6 annotations for the 3 gene scores were significantly enriched when meta-analyzed across 11 blood and autoimmune traits. However, SNP annotations corresponding to `Robospan` and `pRobospan` gene sets showed higher magnitude of enrichment than `Corspan` genes (Figure 4 and Table S4). More importantly, 2 `Robospan`, 2 `pRobospan` and 0 `Corspan` annotations showed significant *τ*^⋆^ conditional on the baseline-LD annotations after Bonferonni correction (Figure 4 and Table S4). If we restrict our analysis to only the 5 autoimmune traits, 2 `Robospan`, 0 `pRobospan` and 0 `Corspan` SNP annotations showed unique signal (Table S5). Even when these annotations were modeled jointly with specifically expressed genes in Whole Blood^38^ (SEG-Blood) and subjected to forward stepwise elimination similar to ref.^41,45,7^, 1 `Robospan` annotation (100kb) still remains significantly informative, suggesting unique disease information over SEG-Blood genes. The same annotation also remains significant in a joint model of just the `Robospan`, `Corspan` and `pRobospan` annotations.

**Figure 4.**
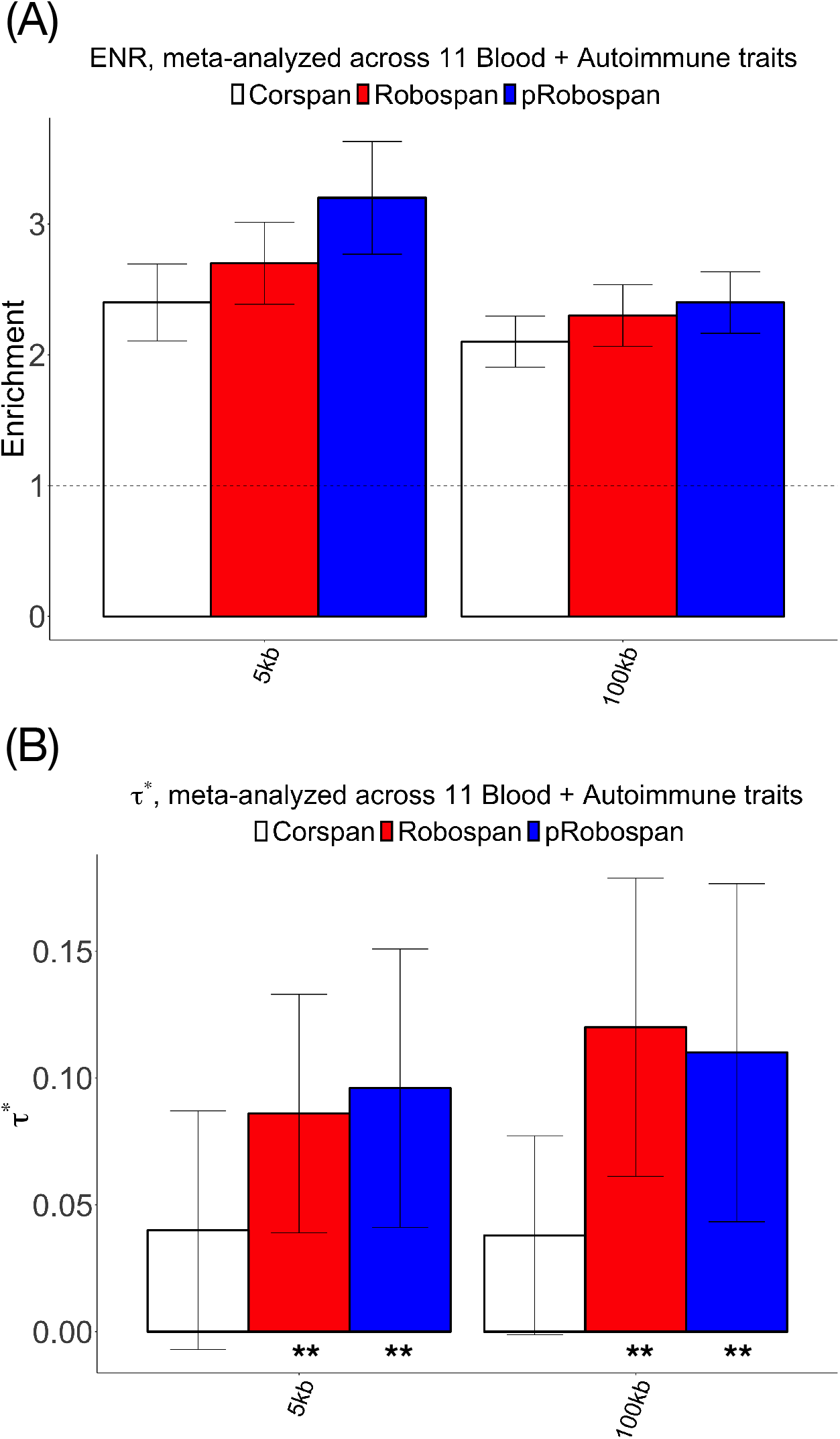
Disease informativeness of 5kb and 100kb SNP annotations for Corspan, Robospan and pRobospan gene sets: (A) Heritability enrichment, conditional on baseline-LD model (v2.1). The base enrichment level is 1. (B) Standardized effect size (*τ*^⋆^) conditional on baseline-LD model for `Corspan` (left column, white), `Robospan` (middle column, red) and `pRobospan` (right column, blue) gene sets. Results are meta-analyzed across 11 blood and autoimmune traits. ** denotes annotations that are significant after Bonferonni correction (*P* < 0.05/8) where 8 is the total number of SNP annotations tested. Error bars denote 95% confidence intervals. Numerical results are reported in Table S4.

We conclude that `Robospan` and `pRobospan` gene sets constructed from `Robocov` correlation and partial correlation estimator show higher enrichment and disease informative signal compared to the `Corspan` gene set constructed similarly from the standard correlation estimator. Additionally, the `Robospan` gene set shows unique disease information (*τ*^⋆^) conditional on the specifically expressed gene in blood; this shows that the study of correlation structure of gene expression across tissues adds value over study of expression data alone.

## 4 Discussion

In this paper we present `Robocov`—a novel convex optimization-based framework for estimating a sparse covariance (correlation) and inverse covariance (partial correlation) matrix, given a data matrix with missing entries. Our approach does not rely on missing data imputation and hence mitigates the possible shortcomings of a sub-optimal imputation procedure (e.g., based on a low-rank assumption). Instead, `Robocov` directly estimates the correlation matrix or partial correlation matrix of interest via a regularized loss minimization framework. Since `Robocov` is a stand-alone generic tool that can be applied to any data with missing entries, it can be used as an exploratory tool for other missing-data related problems.

We have assessed the significance of our proposed `Robocov` framework over standard methods from a methodological, biological and disease analysis perspective. `Robocov` leads to sparse estimates and has a lower false positive rate compared to other competing methods. `Robocov` estimator is visually more interpretable and less cluttered and captures more robust biological signal. In terms of disease informativeness, Robospan and `pRobospan` gene sets, generated from the `Robocov` estimated correlation and partial correlation matrices, perform considerably better than the analogous `Corspan` gene set from standard correlation estimator.

There are several directions for future research. First, `Robocov` may be improved by incorporating the structure of missing values by using additional covariates. In fact, for the GTEx expression data, there may be structured missing-ness driven by post-mortem donor metadata, such as cause of death, age, gender etc. Second, we restrict our study to gene expression data; alternatively, one could have considered transcript expression data. However, accounting for patterns of transcriptional diversity for a particular gene will require more involved modeling assumptions. `Robocov` can also be used to as an ingredient in item response models for large scale data, as in UK Biobank, where there are extensive amounts of missing entries in the response phenotype data^46,47,48^.

`Robocov` is implemented as an R package hosted on Github (https://github.com/kkdey/Robocov).

## URLs

- `Robocov` software https://github.com/kkdey/Robocov
- GTEx v6 data analysis https://github.com/kkdey/Robocov-pages
- List of all genes https://github.com/kkdey/Robocov-pages/tree/master/data/gene_names_GTEx.txt
- Gene Sets https://github.com/kkdey/Robocov-pages/tree/master/Gene_Sets
- Pathway enrichments https://github.com/kkdey/Robocov-pages/tree/master/Pathways
- Annotations analyzed in this study: https://github.com/kkdey/Robocov-pages/tree/master/Annotations
- Baseline-LD annotations: https://data.broadinstitute.org/alkesgroup/LDSCORE/
- 1000 Genomes Project Phase 3 data: ftp://ftp.1000genomes.ebi.ac.uk/vol1/ftp/release/20130502
- UK Biobank summary statistics: https://data.broadinstitute.org/alkesgroup/UKBB/
- Other summary statistics: https://data.broadinstitute.org/alkesgroup/sumstats_formatted/

## Acknowledgements

We would like to thank Alkes L. Price, Bryce van de Geijn and Rajarshi Mukherjee for helpful comments. Rahul Mazumder was partially supported by the Office of Naval Research ONR-N000141512342, ONR-N000141812298 (Young Investigator Award), the National Science Foundation (NSF-IIS-1718258) and IBM. This research was conducted using the UK Biobank Resource under application 16549.

---

**Table S1.** Predictive comparison of CorShrink, Robocov and sample correlation estimators for a GTEx gene. We report Mean Absolute Deviation (MAD) and Root Mean Squared Deviation (RMSD) metrics between an estimator (e.g., sample correlation matrix, `CorShrink` and `Robocov`) computed on the gene expression data (GTEx project) for half of the individuals (training set) and the sample correlation matrix computed from other half (testing set) of all individuals. Results are averaged over 30 such different training/testing data-splits with the standard errors reported in brackets.

| Method | MAD | RMSE |
| --- | --- | --- |
| Sample-Est | 0.30 (0.01) | 0.47 (0.02) |
| CorShrink | 0.24 (0.01) | 0.35 (0.01) |
| Robocov | 0.25 (0.01) | 0.36 (0.01) |

**Table S2.** Pathway enrichment analysis of `Robospan`, `pRobospan` and `Corspan` genes. Pathway enrichment is performed using the ConsensusPathDB database^36,37^. Only the top 5 non-redundant and statistically significant (q-value < 0.05) pathways for a gene set are reported.

| Gene Set | Top pathways |
| --- | --- |
| **Robospan** | Interferon signaling (1.1e-18), Immune system (3.1e-08), HSF1 activation (1.1e-07), Antigen processing and presentation (2.4e-07), Allograft rejection (1.5e-06) |
| **pRobospan** | Immune system (2.7e-21), Interferon signaling (3.4e-15), Innate immune system (5.1e-12), TNF signaling pathway (7.9e-11), Neutrophil degranulation (2.0e-10) |
| **Corspan** | Interferon signaling (1.1e-17), Immune system (1.6e-07), Antigen processing and presentation (1.2e-05), HSF1 activation (4.1e-05), Neutrophil degranulation (1e-04), |

**Table S3.** List of all blood-related traits: List of 11 blood and autoimmune traits (5 blood traits and 6 autoimmune traits) analyzed in this paper.

| Annotation | Traits |
| --- | --- |
| Blood traits | Red blood Cell Distribution Width (UKBB<sup>49</sup>), Red blood Cell Count (UKBB<sup>49</sup>), White blood Cell Count (UKBB<sup>49</sup>), Platelet Count (UKBB<sup>49</sup>), Eosinophil Count (UKBB<sup>49</sup>) |
| Immune traits | Ulcerative Colitis<sup>50</sup>, Rheumatoid Arthritis<sup>51</sup>, Celiac<sup>52</sup>, Lupus<sup>53</sup>, Crohn's disease<sup>50</sup>, Auto Immune and Inflammatory traits |

**Table S4.** S-LDSC results for SNP annotations corresponding to Robospan, pRobospan, Corspan and SEG-Blood gene sets for blood and autoimmune traits.: Standardized Effect sizes (*τ*^⋆^) and Enrichment (E) of 8 SNP annotations corresponding to 4 gene sets (`Robospan`, `pRobospan`, `Corspan` and SEG-Blood^38^) and 2 SNP annotations corresponding to 5kb and 100kb window based SNP-to-gene linking strategies for each gene set. Results for all annotations are conditional on 86 baselineLD-v2.1 annotations. Reports are meta-analyzed across 11 Blood and Autoimmune traits.

| Robospan |  |  |  |  |  |  |
| --- | --- | --- | --- | --- | --- | --- |
| | $\tau^*$ | $se(\tau^*)$ | $p(\tau^*)$ | $E$ | $se(E)$ | $p(E)$ |
| 5kb (2.6%) | 0.086 | 0.024 | 0.00048 | 2.7 | 0.16 | 1.5e-07 |
| 100kb (10%) | 0.12 | 0.03 | 7.9e-05 | 2.3 | 0.12 | 2e-09 |
| pRobospan |  |  |  |  |  |  |
| | $\tau^*$ | $se(\tau^*)$ | $p(\tau^*)$ | $E$ | $se(E)$ | $p(E)$ |
| 5kb (2.3%) | 0.096 | 0.028 | 0.00057 | 3.2 | 0.22 | 9.3e-08 |
| 100kb (9.9%) | 0.11 | 0.034 | 0.0011 | 2.4 | 0.12 | 5.5e-09 |
| Corspan |  |  |  |  |  |  |
| | $\tau^*$ | $se(\tau^*)$ | $p(\tau^*)$ | $E$ | $se(E)$ | $p(E)$ |
| 5kb (2.5%) | 0.04 | 0.024 | 0.093 | 2.4 | 0.15 | 4.8e-07 |
| 100kb (9.8%) | 0.038 | 0.02 | 0.059 | 2.1 | 0.1 | 1.7e-08 |
| SEG-Blood |  |  |  |  |  |  |
| | $\tau^*$ | $se(\tau^*)$ | $p(\tau^*)$ | $E$ | $se(E)$ | $p(E)$ |
| 5kb (2.7%) | 0.24 | 0.036 | 7.6e-11 | 3.6 | 0.26 | 8.7e-10 |
| 100kb (10.1%) | 0.21 | 0.029 | 1.3e-13 | 2.4 | 0.095 | 2.2e-10 |

**Table S5.** S-LDSC results for SNP annotations corresponding to Robospan, pRobospan, Corspan and SEG-Blood gene sets for 6 autoimmune traits.: Standardized Effect sizes (*τ*^⋆^) and Enrichment (E) of 8 SNP annotations corresponding to 4 gene sets (`Robospan`, `pRobospan`, `Corspan` and SEG-Blood^38^) and 2 SNP annotations corresponding to 5kb and 100kb window based SNP-to-gene linking strategies for each gene set. Results for all annotations are conditional on 86 baselineLD-v2.1 annotations. Reports are meta-analyzed across 6 Autoimmune traits.

| Robospan |  |  |  |  |  |  |
| --- | --- | --- | --- | --- | --- | --- |
| | $\tau^*$ | $se(\tau^*)$ | $p(\tau^*)$ | $E$ | $se(E)$ | $p(E)$ |
| 5kb (2.6%) | 0.12 | 0.036 | 0.00074 | 2.7 | 0.25 | 5.2e-05 |
| 100kb (10%) | 0.14 | 0.049 | 0.0051 | 2.3 | 0.19 | 2.1e-06 |
| pRobospan |  |  |  |  |  |  |
| | $\tau^*$ | $se(\tau^*)$ | $p(\tau^*)$ | $E$ | $se(E)$ | $p(E)$ |
| 5kb (2.3%) | 0.1 | 0.043 | 0.016 | 3.3 | 0.39 | 0.00012 |
| 100kb (9.9%) | 0.11 | 0.059 | 0.061 | 2.5 | 0.24 | 1e-05 |
| Corspan |  |  |  |  |  |  |
| | $\tau^*$ | $se(\tau^*)$ | $p(\tau^*)$ | $E$ | $se(E)$ | $p(E)$ |
| 5kb (2.5%) | 0.08 | 0.035 | 0.021 | 2.5 | 0.23 | 0.00011 |
| 100kb (9.8%) | 0.037 | 0.035 | 0.28 | 2 | 0.15 | 7.9e-06 |
| SEG-Blood |  |  |  |  |  |  |
| | $\tau^*$ | $se(\tau^*)$ | $p(\tau^*)$ | $E$ | $se(E)$ | $p(E)$ |
| 5kb (2.7%) | 0.33 | 0.042 | 5.5e-15 | 4.2 | 0.25 | 8.4e-06 |
| 100kb (10.1%) | 0.3 | 0.036 | 8.8e-17 | 2.7 | 0.11 | 7.4e-06 |

**Table S6.** **Joint S-LDSC results for annotations corresponding to `Robospan`, `pRobospan`, `Corspan` and SEG-Blood gene sets.**: Standardized Effect sizes (*τ*^⋆^) and Enrichment (E) of SNP annotations that are significant when all annotations from Table S4 are modeled jointly and subjected to forward stepwise elimination. 2 annotations survive in the resulting joint model. The analysis is conditional on 86 baselineLD-v2.1 annotations. Reports are meta-analyzed across 11 Blood and Autoimmune traits.

| | $\tau^*$ | $\text{se}(\tau^*)$ | $\text{p}(\tau^*)$ | $E$ | $\text{se}(E)$ | $\text{p}(E)$ |
| --- | --- | --- | --- | --- | --- | --- |
| Robospan (100kb) | 0.11 | 0.03 | 0.0001 | 2.3 | 0.12 | 1.8e-09 |
| SEG-Blood (100kb) | 0.21 | 0.029 | 2.3e-13 | 2.4 | 0.095 | 2.1e-10 |

## 5 Supplementary Figures

**Figure S1.**
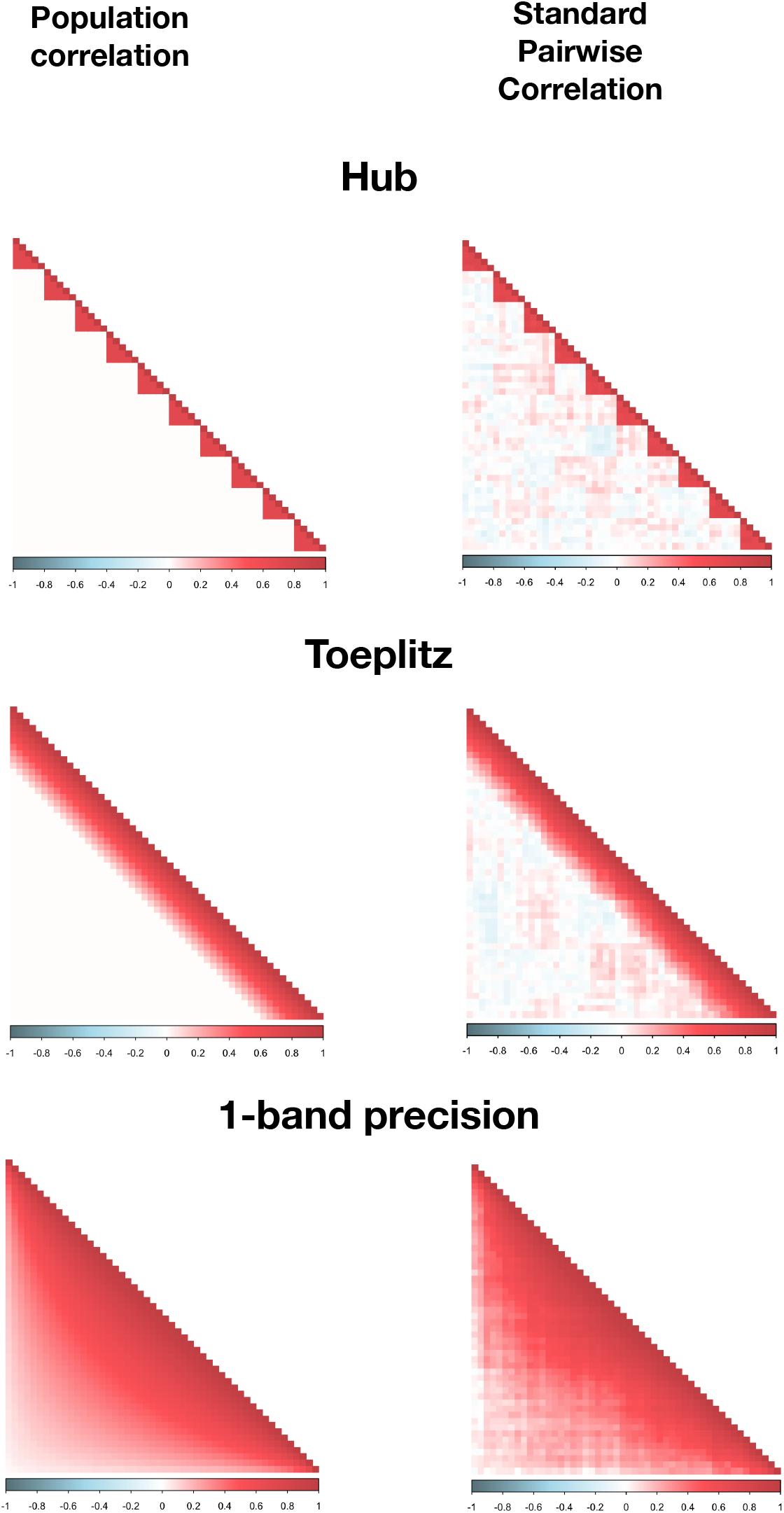
Simulation results of standard pairwise correlation estimator for Hub, Toeplitz and 1-band precision matrices: We applied standard pairwise correlation estimator on data generated from the simulation models from Figure 1-this comprises of Hub, Toeplitz or 1-band precision matrix-based population models with *N*= 500 samples, *P* = 50 features and *π*= 50% proportion of missing data.

**Figure S2.**
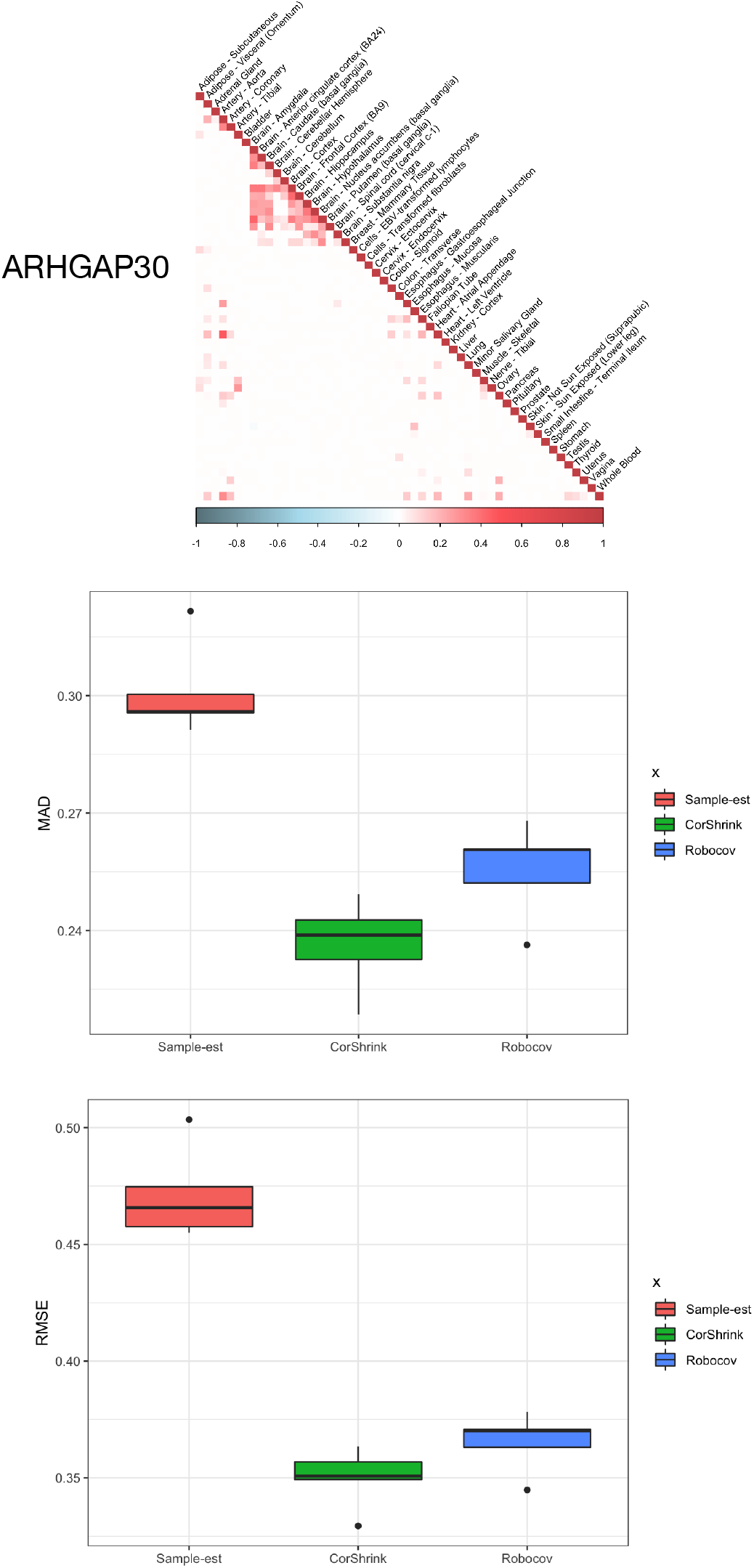
Evaluating the predictive accuracy of Robocov: (Top panel) `Robocov` correlation estimate of the **ARHGAP30** gene. (Middle and bottom panels) We split the data matrix randomly into 2 equal groups. We compare the `Robocov`, `CorShrink` and pairwise sample correlation estimators from one half of the data with the pairwise sample correlation matrix on the other half. We use Median Absolute Deviation (MAD) (middle panel) and Root Mean Squared Error (RMSE) (lower panel) metrics. The results are averaged over 50 such random splits. See Table S1 for a numerical summary.

**Figure S3.**
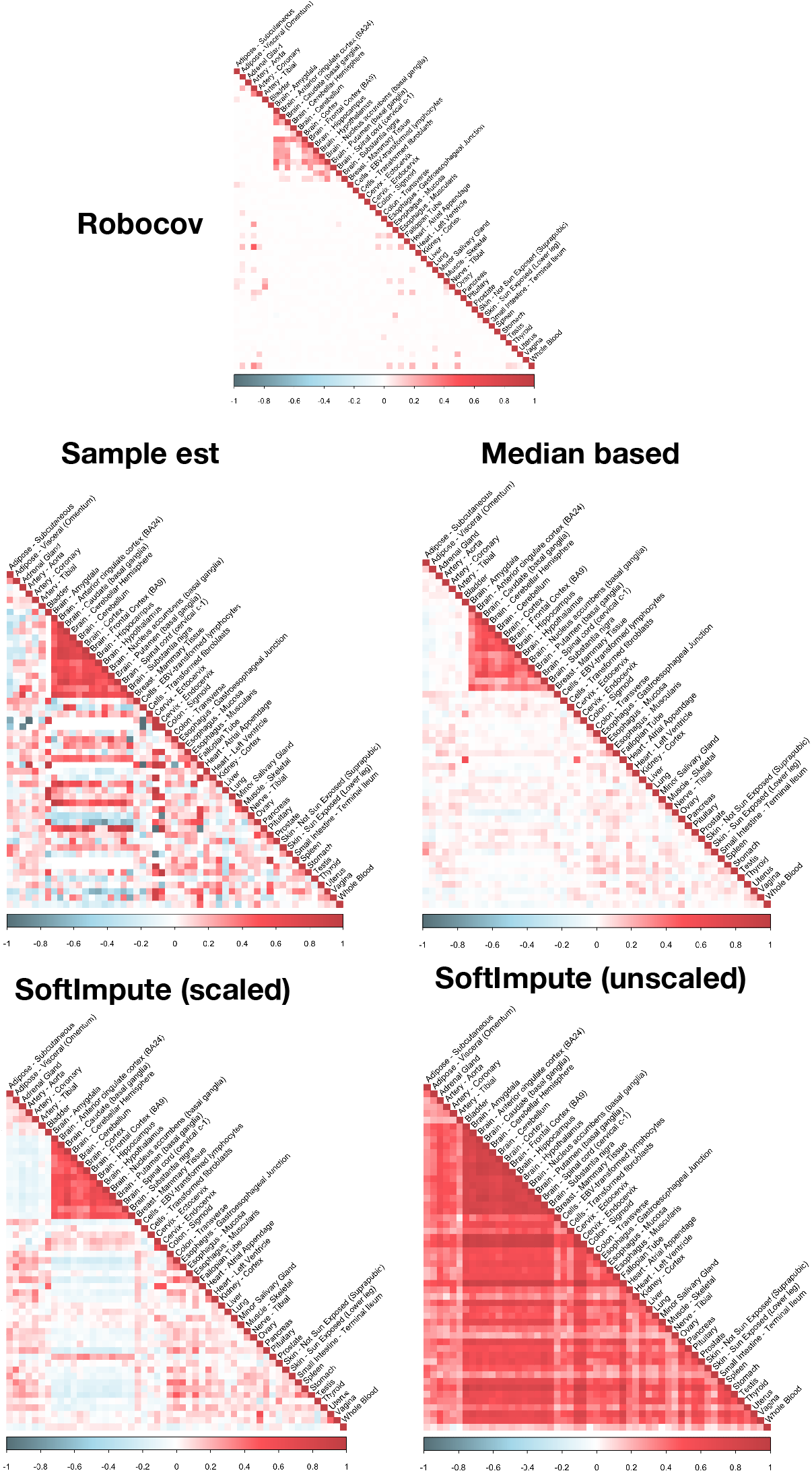
Comparison of the Robocov correlation estimator with correlation estimators based on imputed data: We compare the `Robocov` correlation estimator for the ARHGAP30 gene with four other estimators. They include the standard pairwise sample correlation estimator, the sample correlation matrix computed over data imputed by either a median-based approach (missing entries of a feature replaced by the median of observed entries), the scaled SoftImpute^9^ approach; and an unscaled SoftImpute^9^ approach.

**Figure S4.**
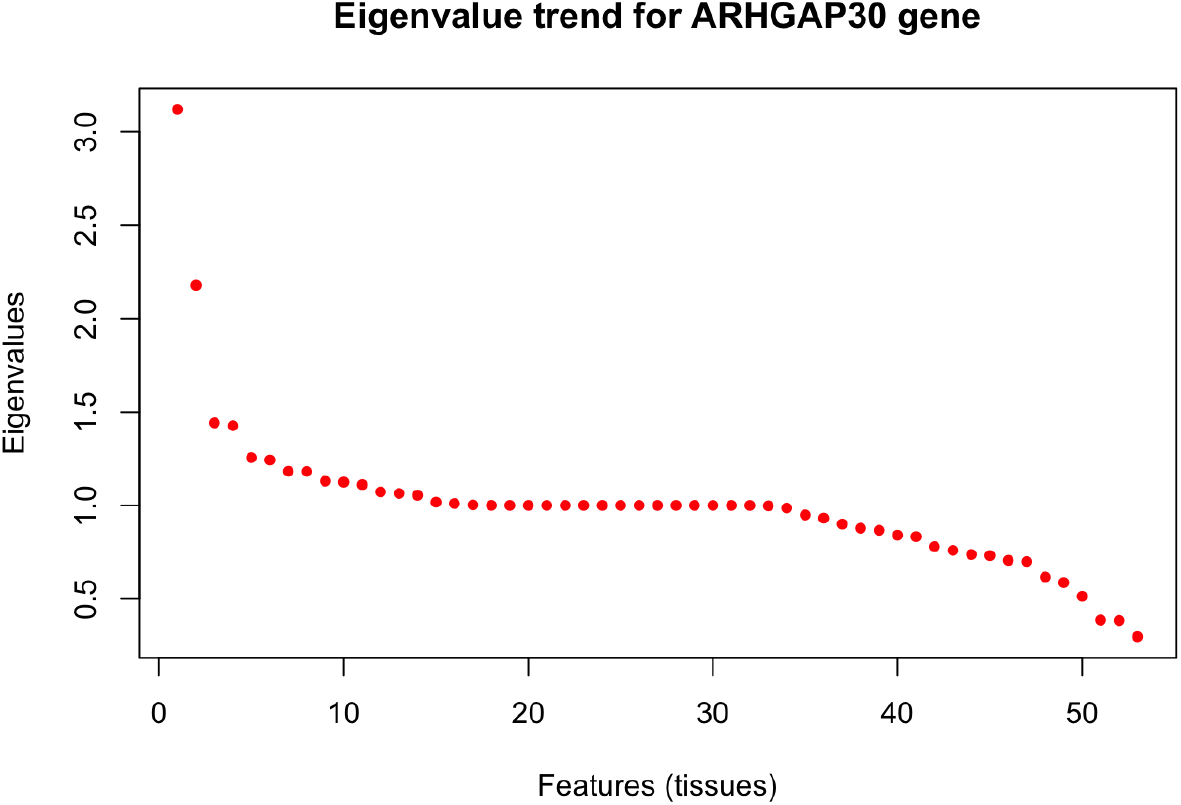
Illustration of high rank for GTEx tissue-tissue correlation matrix: Plot of eigenvalues sorted from highest to lowest in magnitude for tissue-tissue pairwise correlation matrix for a particular gene (ARHGAP30). The eigenvalues do not show any sharp drop close to 0 as one would expect if the matrix allowed a low rank (+noise) structure. This suggests relatively high dimensional structure in the GTEx gene expression data which may explain why a low rank imputation method such as SoftImpute^9^ performs poorly in S3.

**Figure S5.**
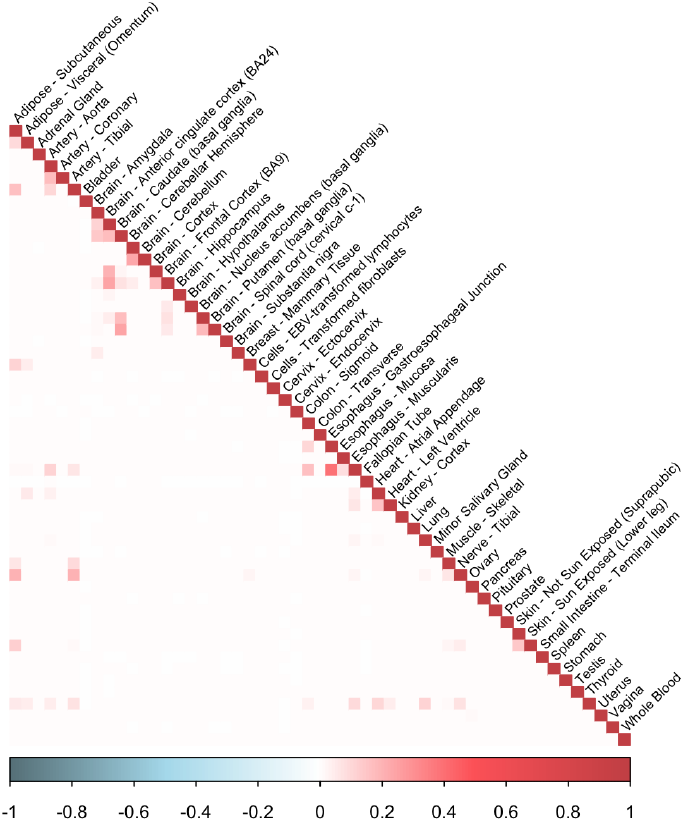
Median of Robocov correlation estimators: We compute median of `Robocov` correlation estimates for each tissue-tissue pair across 16,069 genes studied.

**Figure S6.**
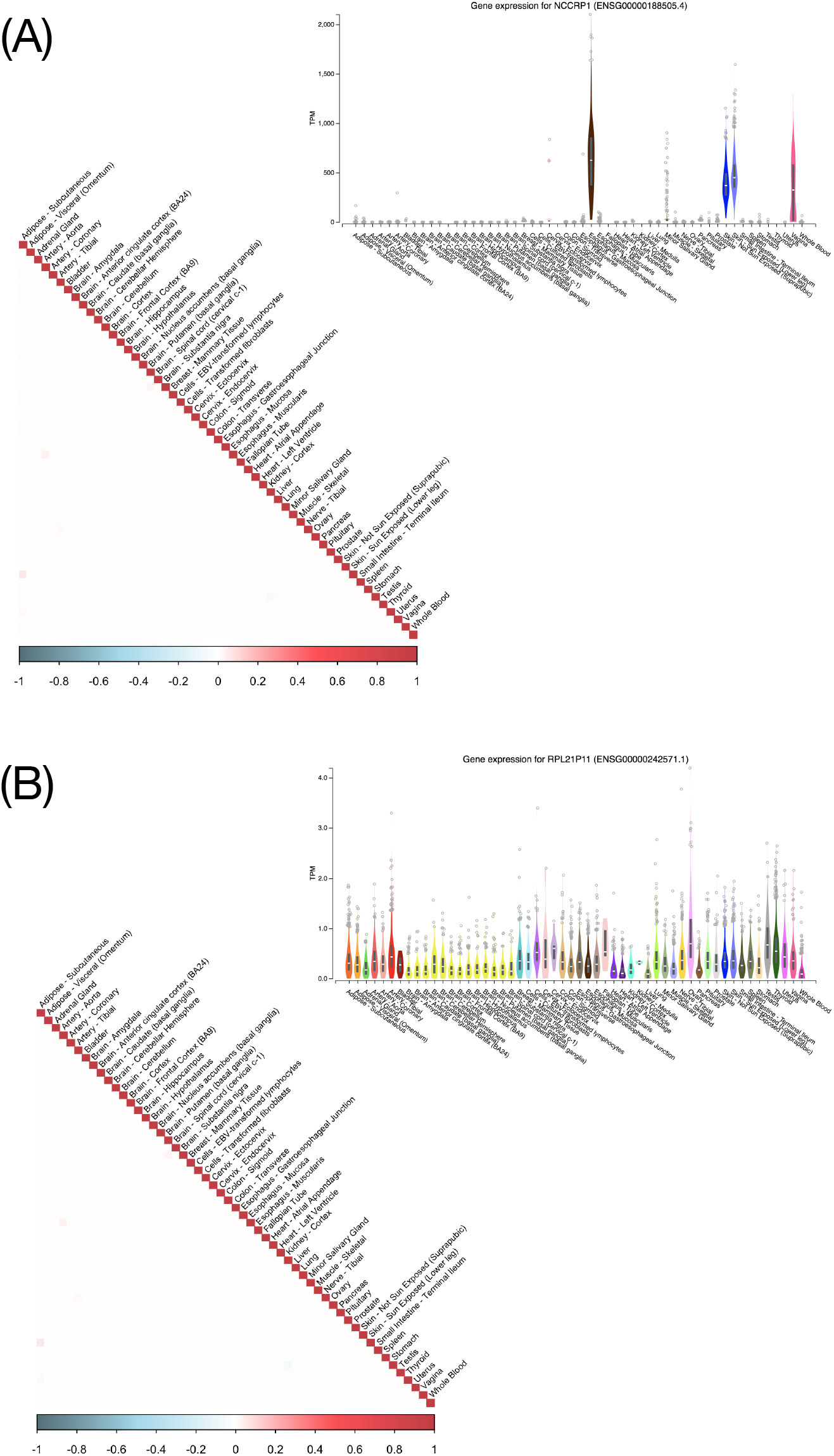
Examples of two genes with low Robospan-score but having distinct expression patterns: Examples of two genes, **NCCRP1** (top) and **RPL21P11** (bottom), both of which have close to 0 average correlation in expression across tissue-pairs but having very distinctive expression profiles. **NCCRP1** has high expression in a few specific tissues including Whole Blood, while **RPL21P11** has uniformly low expression across all tissues. The expression profile plots for the genes have been fetched from the GTEx Portal (https://gtexportal.org/home/).

**Figure S7.**
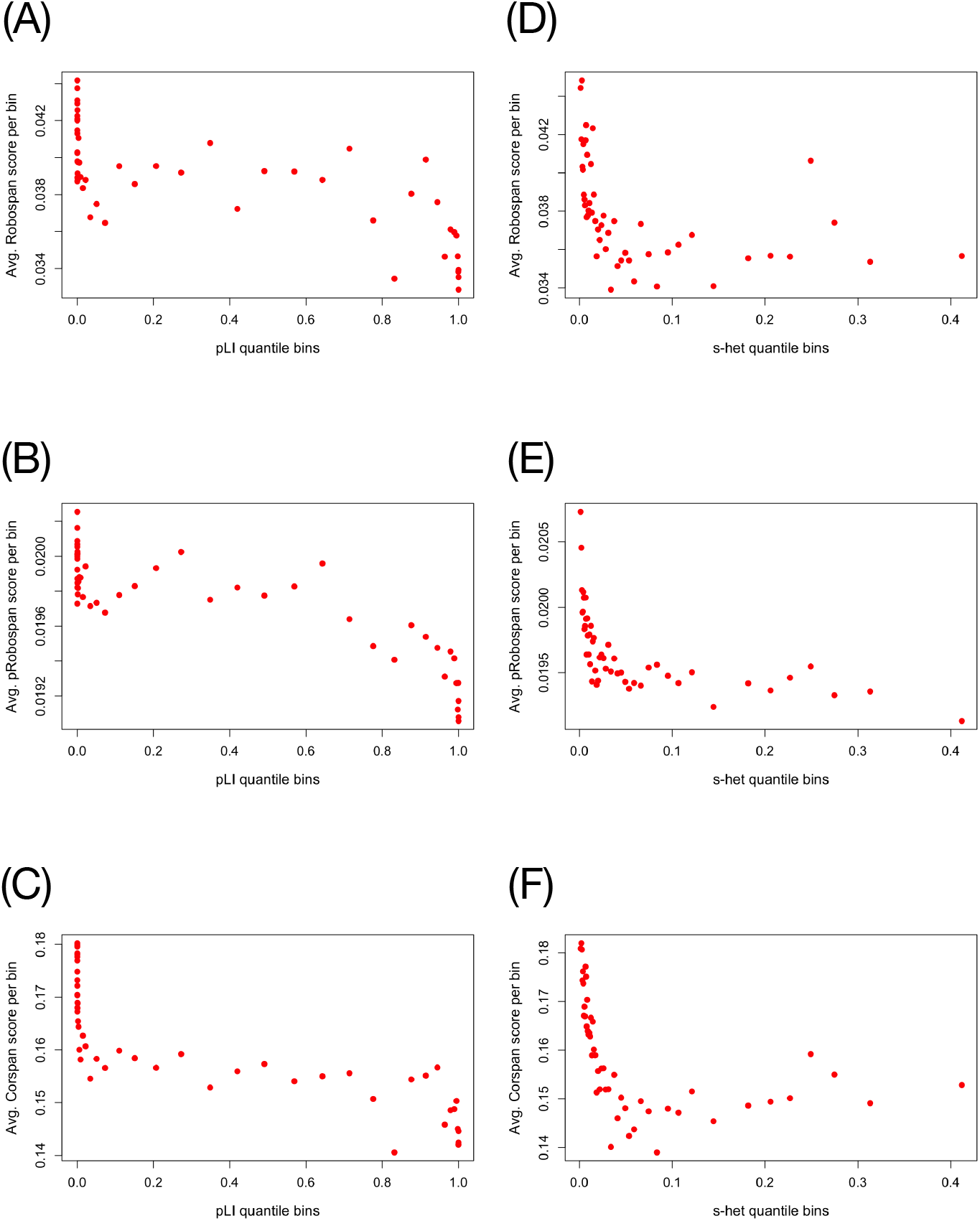
Comparing Robospan-score, pRobospan-score and Corspan-score with pLI and s_het: Comparison of pLI gene score with (A) Robospan-score, (B) `pRobospan`-score and (C) Corspan-score for all genes (See Results section for details). Comparison of s_het gene score with (A) Robospan-score, (B) `pRobospan`-score and (C) Corspan-score for all genes (See Results section for details).

**Figure S8.**
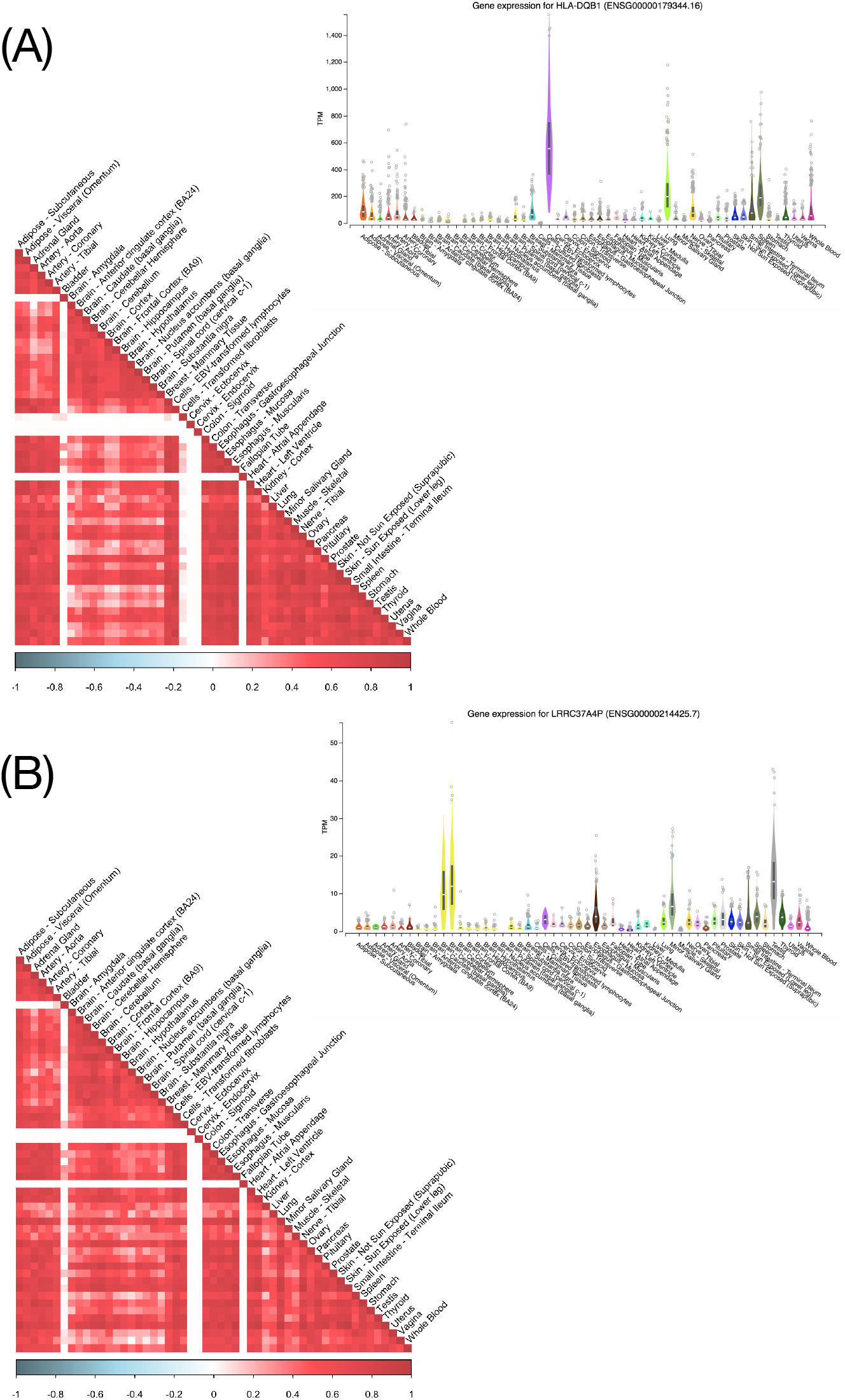
Examples of genes high Robospan-score but not specifically expressed in blood or uniformly expressed across tissues: The two genes are **HLA-DQB1** (top) and **LRRC37A4P** (bottom). **HLA-DQB1** is specifically expressed in lymphocyte cell line which is related to blood. **LRRC37A4P** has highest expression in brain cerebellum and testis. The expression profile plots for the genes have been fetched from the GTEx Portal (https://gtexportal.org/home/).

**Figure S9.**
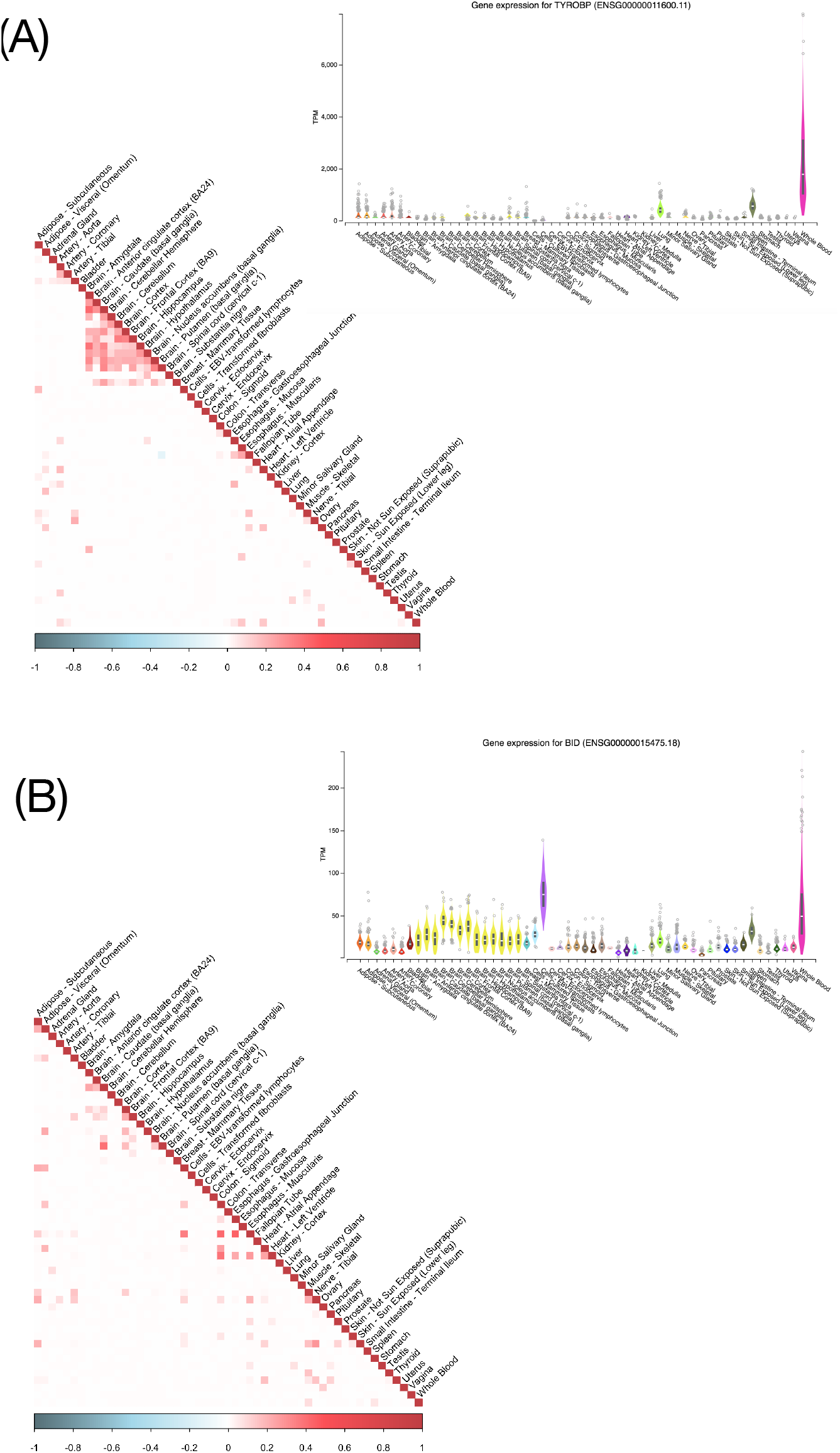
Examples of genes that are specifically expressed in Whole Blood but do not show high Robospan-score: Examples of two genes, **TYROBP** (top) and **BID** (bottom), that show tissue specific expression in Whole blood and are in top 10% specifically expressed genes (SEG) as per ref.^38^, but do not show consistently high expression correlation across tissue pairs and hence, do not have high Robospan score. The expression profile plots for the genes have been fetched from the GTEx Portal (https://gtexportal.org/home/).

### Supplementary Note

#### Fisher Z-score

The population Fisher Z-score^25^ is defined as

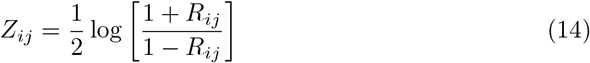

where *R* is the population correlation matrix. The corresponding empirical Fisher Z-score is defined as follows

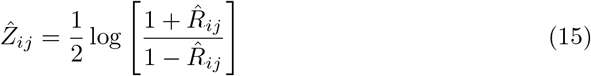

For bivariate normally distributed random variables *X_i_* and *X_j_*, the empirical Fisher Z-score *Ẑ_ij_* (based on *n_ij_*-many samples) is normally distributed given the population counterpart *Z_ij_*^26^:

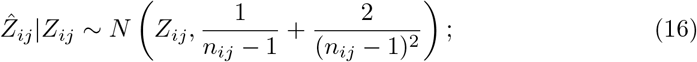

and the Z-scores are conditionally independent. Dey and Stephens^7^ assume an adaptive shrinkage prior on the population Fisher Z-scores for each pair of variables. Here we use property (16) in the context of directly estimating Σ or Ω with an *ℓ*_1_-norm penalty.

#### Derivation of C

Here we show how we derive the analytical form of the upper bound C in (5) appearing in Problem (4).

##### Lemma 1.

*Let* 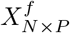 *be the fully observed version of the data matrix X*; *and let every sample* 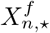 *follow a Multivariate Gaussian distribution with covariance matrix* Σ *and correlation matrix R. The samples are independent. Then, for any fixed ϵ* > 0 *and for sufficiently large n_ij_, there exists a* 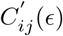 *such that*

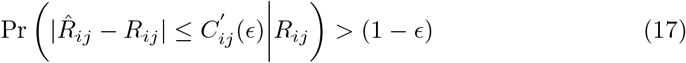

*where*

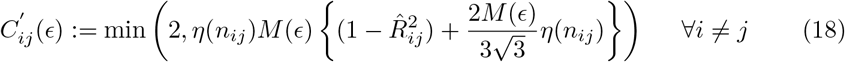

*and*

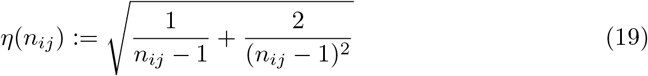

*and M*(*ϵ*) *is a sufficiently large finite number*.

##### Corollary 1.

*For ϵ* = 0.001, *M*(*ϵ*) *can be taken to be* 3 *in Lemma 1. Then*

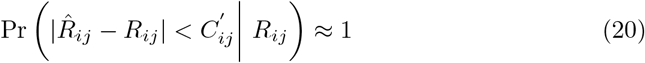

*where*

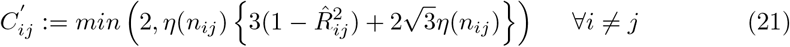

If *n_i_* and *n_j_* are sufficiently large, in which case 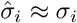 and 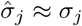, then Corollary 1 leads to the following probability inequality for the pairwise sample covariance:

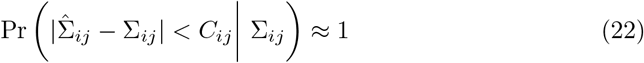

where

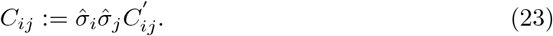

#### Proof of Lemma 1and Corollary 1

If a random variable *W* ~ *N* (0, 1), then for any small *ϵ* > 0, we can get a number *M*(*ϵ*) such that

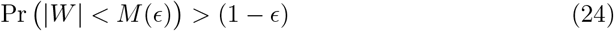

Using (16) and (24), we have

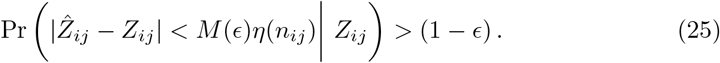

The estimated and population correlations 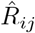 and *R_ij_* (respectively) can be written in terms of the Z-scores using (14) as follows:

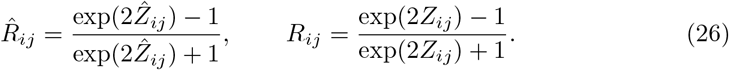

Applying a Taylor series expansion to *R_ij_* as a function of *Z_ij_* around *Ẑ_j_*, we get:

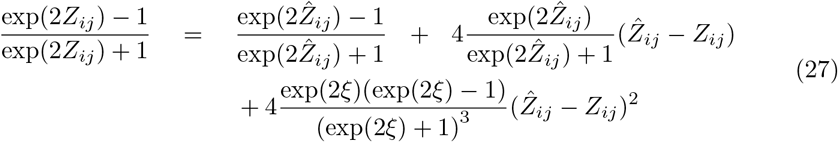

where *ξ* is a value between *Z_ij_* and *Ẑ_ij_*. We can place an upper bound on the coefficient of the last term in (27):

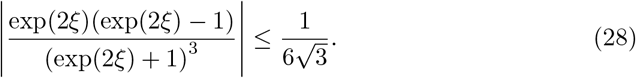

Using Equations (26), (27) and (28), we can write

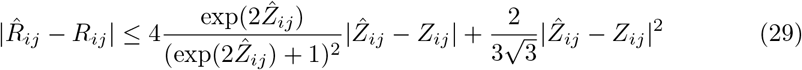

Using the definition of *Ẑ_j_* in Equation (15), we get

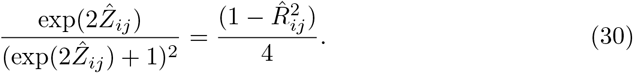

Using the above expression in (29), we get:

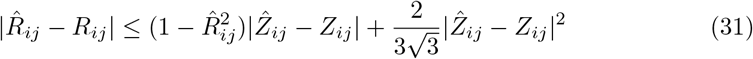

Using (25) and (31), we have:

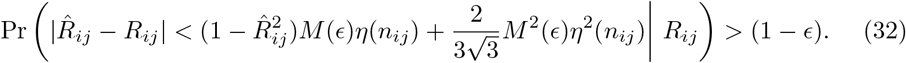

Since, 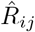 and *R_ij_* are both correlation terms, they lie between −1 and +1 and hence with probability one:

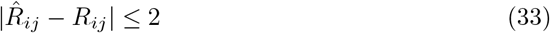

Combining Equations (31) and (33), we get

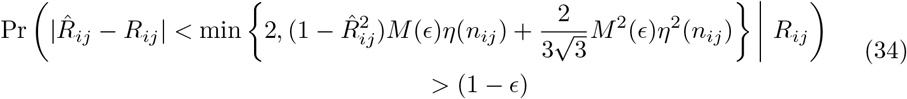

which completes the proof of Lemma 1.

In (24), if we choose *ϵ* = 0.001, we have *M*(*ϵ*) ≈ 3—hence, (34) leads to:

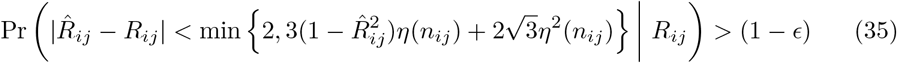

which proves Corollary 1. Usually this result holds good^26^ for any *n_ij_* > 3. If however *n_ij_* → ∞ for all (*i, j*) pairs, then the bound on 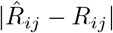 in (34) approaches 0 and 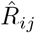 would be close to *R_ij_*.

### A General Likelihood Framework for Robocov Covariance Matrix Estimation

We propose a generalization of the `Robocov` covariance matrix estimation framework presented in Section 2.1. We present a family of loss functions for the regularized criterion (7)— the loss function presented here is directly motivated by the Fisher’s Z-score framework discussed above, but differs from that appearing in Section 2.1.

Recall that the estimators in Section 2.1 are special cases of the following regularized loss minimization framework:

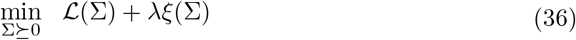

where 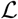 is the data fidelity function and *ξ* is the penalty function. In Section 2.1, we consider an *ℓ*_1_-penalty on the entries of Σ — i.e., 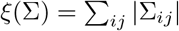. We present below (See (37)) a convex quadratic loss function 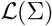. While this differs from the loss function considered in (4), in practice, the performances of these two estimators were found to be similar (at least on the datasets we experimented on).

To derive the loss function, we make use of Lemma 2 — which presents the (conditional) mean and variance of 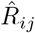 (given *R_ij_*). This leads to a loss function of the form:

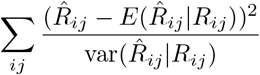

Using the expressions for conditional mean/variances from Lemma 2 (see below), in the above expression, we get:

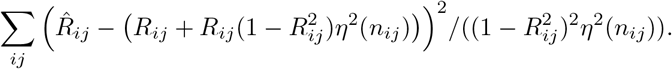

We set 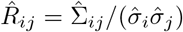 above, and obtain

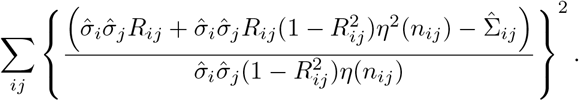

The loss function above is a highly nonconvex function in *R_ij_* or Σ_*ij*_. To this end, we approximate the above by replacing some unknown population quantities by their sample analogues. This results in a loss function:

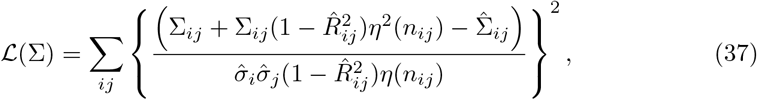

which is convex in Σ. In words, 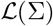 above, is a measure of how close Σ*_ij_*s are to the pairwise covariance terms 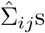—this critically depends upon the number of observed samples *n_ij_* for every pair (*i,j*).

We now present Lemma 2 and its proof:

#### Lemma 2.

*Assume that all conditions of Lemma 1 hold. If n_ij_ is large so that* 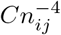 *is negligible for a constant C, we have:*

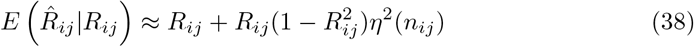

*and*

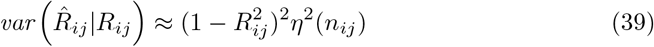

*where η*(*n_ij_*) *is as described in* (19).

#### Proof of Lemma 2

We re-write 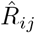 as a function of the Fisher Z-score

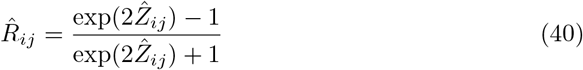

We then expand 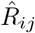 as a function of *Ẑ_ij_* around the population Fisher Z-score *Z_ij_* using the 2nd order Taylor series expansion as follows:

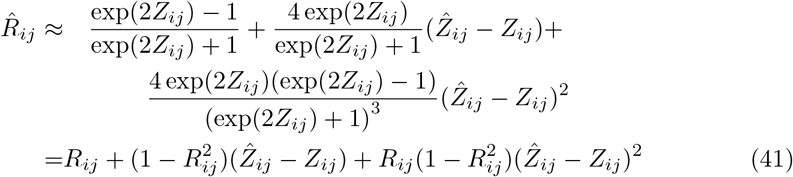

Using the fact that *E*(*Ẑ_ij_*|*R_ij_*) = *E*(*Ẑ_ij_*|*Z_ij_*) = *Z_ij_*, we get from (41)

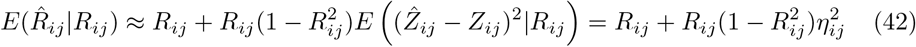

and

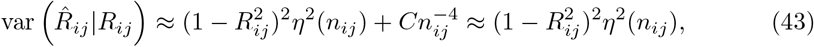

where (43) makes use of the fact that 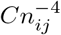 is negligible as per the condition of Lemma 2; and the cross (covariance) term vanishes as it is the third moment of a Gaussian with mean zero.

#### Derivation of D in (11)

Here we discuss how we derive the analytical form of *D* in (11) in the optimization framework in (10).

Let 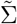 be the sample covariance matrix of *X^f^* (i.e., the fully observed version of *X*) We implicitly assume that the perturbation amount Δ is such that 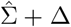 is a good approximation to the unobserved 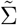. That is,

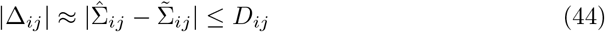

We can write

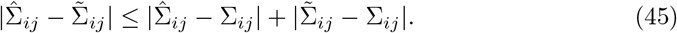

We propose bounds on each of the two terms on the right using our results from the Robocov covariance matrix section. We know that the first term would be bounded by *C_ij_* from Corollary 1. Note that 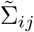 is an instance of 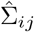 when *n_ij_* = *N* – i.e., all samples are observed. Hence, the bound will be similar to *C_ij_* but with *n_ij_* replaced by *N*. We therefore define

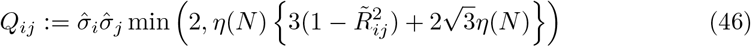

where 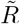 is the correlation matrix corresponding to 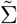.

When *N* is reasonably large, 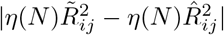 is very small since both 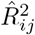 and 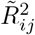 are bounded between 0 and 1 and *η*(*N*) → 0 as *N* → ∞.

Therefore we can effectively replace *Q_ij_* by 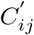 defined as:

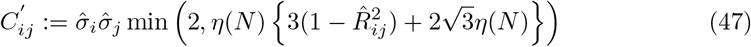

This provides a justification for the choice of *D* appearing in (11).

### Arriving at the Robocov inverse covariance estimator in Section 2.2

Here we explain how the min-max optimization problem in (10) leads to the optimization problem in (12).

To this end, note that:

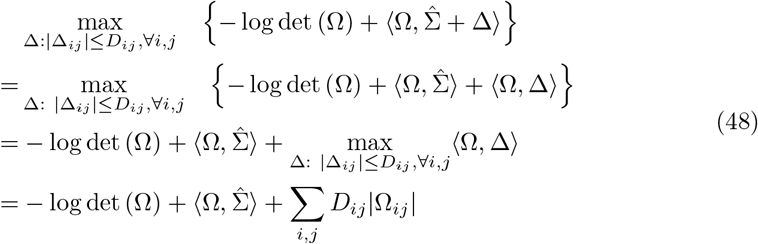

where, the last line follows by noting that

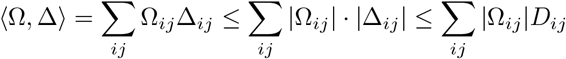

and an equality above holds when Δ_*ij*_ = sign(Ω_*ij*_)|*D_ij_*| for all *i,j*, Using (48), Problem (10) becomes:

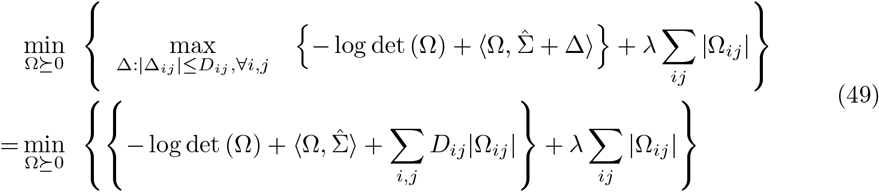

which is the formulation appearing in (12).

### Simulation settings

The parameter models for the simulated population models in Figure 1 are as follows.

- **Hub:** The hub matrix population model for both Figure 1 and Table 1 comprised of correlation blocks of size 5. Each block had all off-diagonal entries equal to 0.7.
- **Toeplitz:** The Toeplitz matrix population model A in Figure 1 had entries of the form *A_ij_* = max {0,1 — 0.1 * |*i* – *j*|}.
- **1-band precision:** The 1-band precision matrix population model in Figure 1 is of the form *A*_*i,i*+1_ = 0.5 an *A_i,j_* = 0 for *j* ≠ *i,i* +1 for each feature *i*.

### Performance metrics

Three performance metrics were used to compare different correlation and partial correlation estimators for different simulation settings (Table 1). They include

- **FP2: False Positive 2-norm**: Euclidean distance of the estimated correlation or partial correlation values for feature pairs with population correlation or partial correlation equal to 0.
- **FPR: False Positive Rate**: The proportion of feature pairs with population correlation (partial correlation) equal to 0 that have estimated correlation (partial correlation) greater than 0.1.
- **FNR: False Negative Rate**: The proportion of feature pairs with population correlation (partial correlation) greater than 0.1 that have estimated correlation (partial correlation) less than 0.01.

### Stratified LD-score regression

Stratified LD score regression (S-LDSC) is a method that assesses the contribution of a genomic annotation to disease and complex trait heritability^40,43^. S-LDSC assumes that the per-SNP heritability or variance of effect size (of standardized genotype on trait) of each SNP is equal to the linear contribution of each annotation

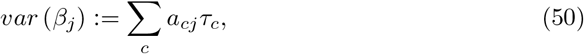

where *a_cj_* is the value of annotation *c* for SNP *j*, where *a_cj_* is binary in our case, and *τ_c_* is the contribution of annotation *c* to per-SNP heritability conditioned on other annotations. S-LDSC estimates the *τ_c_* for each annotation using the following equation

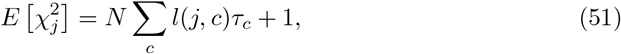

where 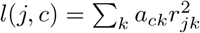 is the *stratified LD score* of SNP *j* with respect to annotation *c* and *r_jk_* is the genotypic correlation between SNPs *j* and *k* computed using data from 1000 Genomes Project^39^ (see URLs); *N* is the GWAS sample size.

We assess the informativeness of an annotation c using two metrics. The first metric is enrichment (*E_c_*), defined as follows (for binary and probabilistic annotations only):

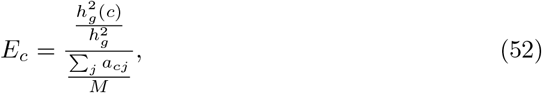

where 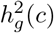 is the heritability explained by the SNPs in annotation *c*, weighted by the annotation values.

The second metric is standardized effect size (*τ*\*) defined as follows (for binary, probabilistic, and continuous-valued annotations):

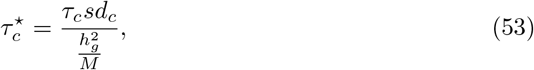

where *sd_c_* is the standard error of annotation *c*, 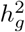 the total SNP heritability and *M* is the total number of SNPs on which this heritability is computed (equal to 5, 961, 159 in our analyses). 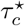 represents the proportionate change in per-SNP heritability associated to a 1 standard deviation increase in the value of the annotation.

## Footnotes

1 Expectation Maximization (EM)^17^ methods are often used for estimation with missing values, but (i) they depend upon probabilistic modeling assumptions on the data; and (ii) they lead to highly nonconvex problems posing computational challenges.

2 Note that the data matrix *X* is a restriction of *X^f^* to the observed entries.

3 If necessary, as a pre-processing step, we remove features so that the condition *n_ij_* < 2 is satisfied for all *i,j*.

4 We get a positive semidefinite (PSD) estimate for Ω even if 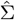 is not PSD. In addition, due to the presence of the logdet in the objective, an optimal solution to (12) will be positive definite (i.e, Ω will have full rank).

